# Genome-resolved house microbiome exhibits location-specific metabolic partitioning, and harbors hosts with clinically relevant antibiotic resistance genes

**DOI:** 10.64898/2026.05.24.727512

**Authors:** Saraswati Awasthi, Rakesh Sharma

**Affiliations:** CSIR-Institute of Genomics and Integrative Biology, New Delhi, India; Academy of Scientific and Innovative Research (AcSIR), Ghaziabad, India

**Keywords:** House microbiome, MAGs, ARGs, metabolism, resistome, clinically relevant

## Abstract

Residential indoor surfaces are recognized as diverse microbial ecosystems, while their genome-based organization and functional repertoires remain understudied. We recovered 2304 metagenome-assembled genomes (MAGs) from shotgun metagenomic sequencing of 10 houses in New Delhi, India. Genome-resolved analysis revealed a highly structured microbial community and substantial unexplored diversity, with 60% Species-level Genome Bins (SGBs) (629/1014) unclassified at the species level. Metabolism reveals a conserved metabolic core, along with spatial functional enrichment: the living area was significantly different from the bathroom and kitchen areas. The prevalent MAG species of the house microbiome, *Paracoccus marcusii*, *Ottowia* sp. 018060485, and *Kocuria palustris*, showed strain-level diversity with no stratification by house, but a subtle location-wise grouping. Potential pathogens, along with a wide range of antimicrobial resistance genes (ARGs), were identified across the MAGs, with 64 ARGs associated with mobile elements. Phylogenomic analysis of *Escherichia coli* MAGs indicated a split between commensal-like fecal lineages and pathotype-associated clusters, like Intestinal Pathogenic *E. coli* (InPEC). These results suggest that residential house microbiomes harbor microbial communities with both diverse metabolic capacity and clinical relevance. Together, these findings establish a reference for future indoor microbiome research and provide a foundation for antimicrobial resistance surveillance and the development of bio-informed building-infrastructures.

## 1. Introduction

Indoor environments constitute complex and dynamic microbial ecosystems. The house microbiome comprises a diverse community of bacteria, archaea, fungi, and viruses^1^. Unlike other environments, residential spaces constitute a unique microbial meeting ground as microbes from outdoor environments (air, water, and soil) converge with additional inputs from human skin shedding and hair^1^. These microbial communities are influenced by factors such as building design, human occupancy, pet presence, materials used, and proximity to green spaces^2,3^. These dynamic ecosystems shape human microbial exposure, modulate immune development, and facilitate the transmission of pathogens and antimicrobial resistance determinants^1,4–7^. With the increase in urbanization, contemporary building designs have isolated residential spaces from outdoor environments, shifting the balance between environmental microbes and human-associated taxa, plausibly altering the health-relevant microbial landscape of homes^8–11^.

The residential microbiome has already been shown to harbor substantial biological diversity. In the Indian house microbiome study, more than 1,400 bacterial species, over 100 viral groups, and 669 antimicrobial resistance gene (ARG) subtypes spanning 22 classes have been reported^12^. In addition, ESKAPE pathogens and opportunistic strains such as *Escherichia coli* have been detected on household floors across different sites, underscoring the need to examine these communities in the context of human health^13^.

Amplicon and short-read metagenomic approaches suffer in terms of strain resolution and the association of function with taxa, which are crucial for understanding the survival of organisms in resource-limited environments^14,15^. Reconstructing metagenome-assembled genomes (MAGs) bridges this gap through metabolic characterization and by linking antimicrobial resistance genes (ARGs) and virulence determinants to their host genomes, allowing assessment of their genomic context^16^. Despite increased knowledge, limited information is available on the functional genomic diversity associated with residential microbes.

In this study, we recovered genomes from shotgun metagenomic data generated from Indian residential samples to assess the functional profiles and clinical implications of indoor microbial communities. A total of 90 samples from 9 locations across 10 houses were analyzed to: (i) expand the genomic representation of indoor residential floor-associated microbes; (ii) elucidate metabolic potential related to carbon, nitrogen, sulfur, and iron cycling; (iii) assess the presence of potential pathogens, prevalence and genomic context of antimicrobial resistance genes in the house microbiome; and (iv) explore the phylogenomic diversity of *Escherichia coli*. Together, these analyses provide genome-resolved insight into the ecological potential and health relevance of residential microbiomes.

## Results

### Recovery of 2304 MAGs from 90 House Microbiome Samples

Homes are not microbiologically uniform: different rooms create distinct habitats that shape which microbes persist and what functions they perform. To understand the microbiome and role of individual member, we used genome-resolved metagenomics from 90 samples across 10 houses in New Delhi, to assemble 2304 MAGs (**Fig. 1a**). These genomes represent medium (n=1642) and near-complete (n=662) MAGs in accordance with the MIMAG^17^ standards and were dereplicated sample-wise. Out of the 662 near-complete bins, 75 were high-quality draft MAGs (**Fig. 1c**). Most of the MAGs were recovered from the bathroom area (n=991) location group followed by the kitchen area (n=743), and then the living area (n=570) (**Fig. 1b**). If looked at different locations of the house, the shower area had the highest MAG counts (n=462) and the bedroom had the least recovery of the MAGs (n=164) (**Suplementary Fig. 1a**). All MAGs were bacterial and none of the archaeal MAGs were recovered. The phyla distribution of the MAGs across the location groups showed dominance of Pseudomonadota across all locations, followed by Actinomycetota. However, the phylum Bacillota remains relatively abundant in the living area as compared with other location groups. Dereplication of the 2,304 MAGs yielded 1,014 SGBs, of which 385 (37.97%) were classified at the species level, and 629 (62.03%) remained unclassified at the species level (**Supplementary Fig. 1c**). At higher ranks, A70_bin.91 (class Actinomycetia) remained unclassified at the order level, while A41_bin.25 (class Alphaproteobacteria, order *UBA1280*) and A86_bin.31 (Vicinamibacterales) remained unclassified at the family level. The unclassified SGBs included several members of the rare biosphere, particularly underexplored environmental lineages such as Patescibacteria, Armatimonadota, Sumerlaeota, Bdellovibrionota, and Myxococcota. The ANI (Average nucleotide identity) vs alignment fraction (AF) scatter plot (**Fig. 1e**) shows the distinctiveness of the recovered genomes relative to the house microbiome data. The resulting genomic landscape revealed a robust sequence-discrete structure, with a high density of MAGs sharing >95% ANI and >0.75 AF, defining clear species-level boundaries for the majority of indoor taxa. However, a large fraction (593) of the genomes have ANI < 95% and AF > 0.5, indicating evolutionary divergence between the co-occurring taxa. These genomes represent phylogenetically divergent or rare members of the household microbial community. The phylogenetic distribution of the house-recovered MAGs reveals a high proportion of genomes belonging to previously uncharacterized species, dispersed across nearly all major clades. Nearly 50% of the house MAGs remain unclassified at the species level (**Fig. 2**, inner color strip). Mapped genomic features across the tree show significant heterogeneity in both genome size (inner bar chart) and GC content (outer bar chart), spanning 0.03-10.18 Mb and 22%-75.6%, respectively.

**Fig. 1.**
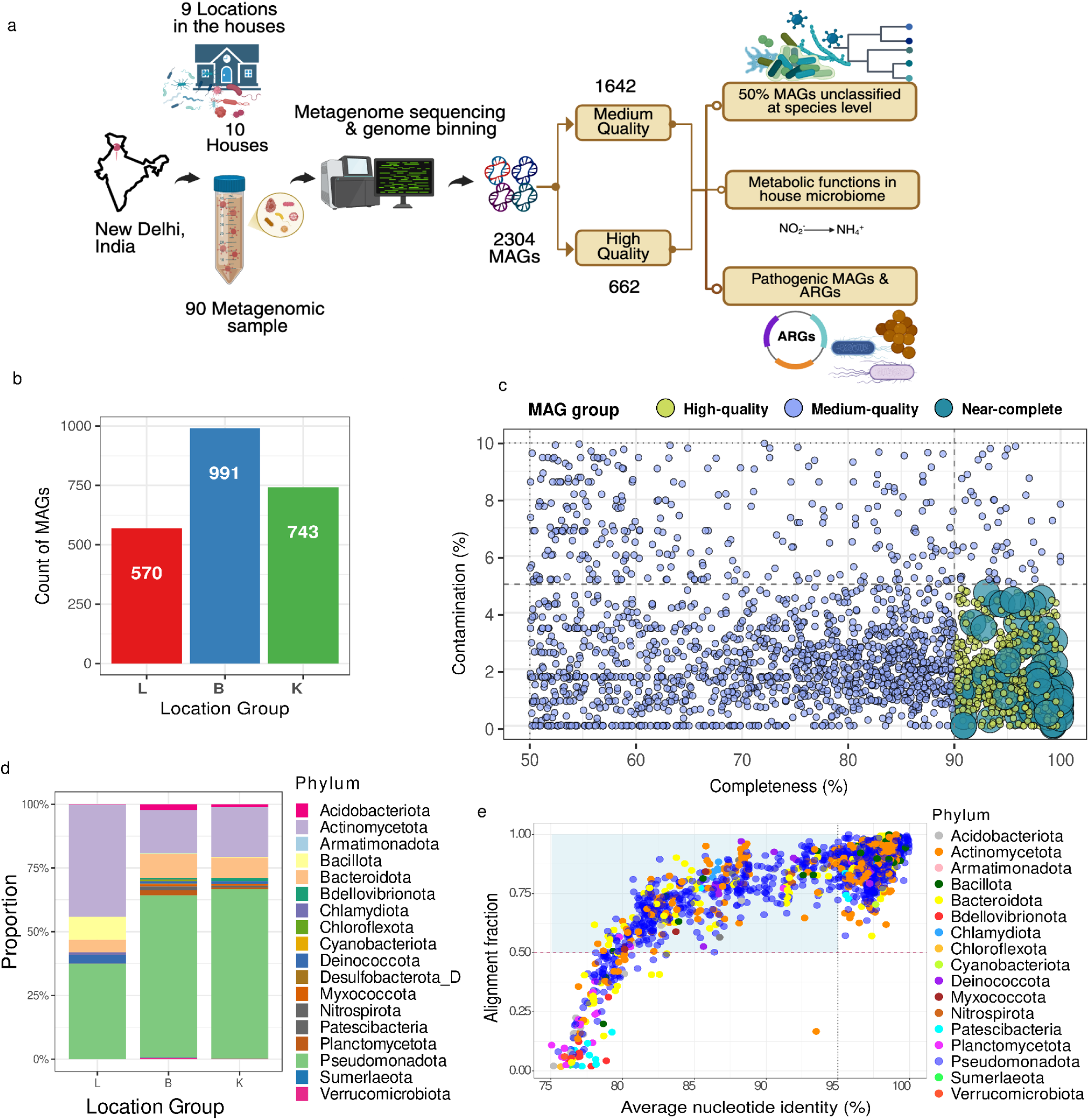
Genome resolve metagenomic of the indoor house microbiome. (a) Overview of the genome-resolved metagenomic workflow, from sampling and sequencing to reconstruction of metagenome-assembled genomes (MAGs) and downstream functional and pathogenicity analyses. (b) MAG recovery across location groups shows clear differences in genome yield, with group B contributing the largest fraction of reconstructed genomes. (c) MAG quality distribution based on completeness and contamination, highlighting a substantial proportion of high-quality (≥90% completeness, ≤5% contamination) and near-complete genomes, alongside a continuum of medium-quality bins. (d) Phylum-level composition of reconstructed MAGs reveals dominance of major bacterial lineages, with location-specific shifts in relative abundance. (e) Scatter plot of Average nucleotide identity (ANI) versus alignment fraction indicates that most MAGs are closely related to known taxa, while a subset with lower similarity suggests the presence of previously uncharacterized genomic diversity.

**Fig. 2.**
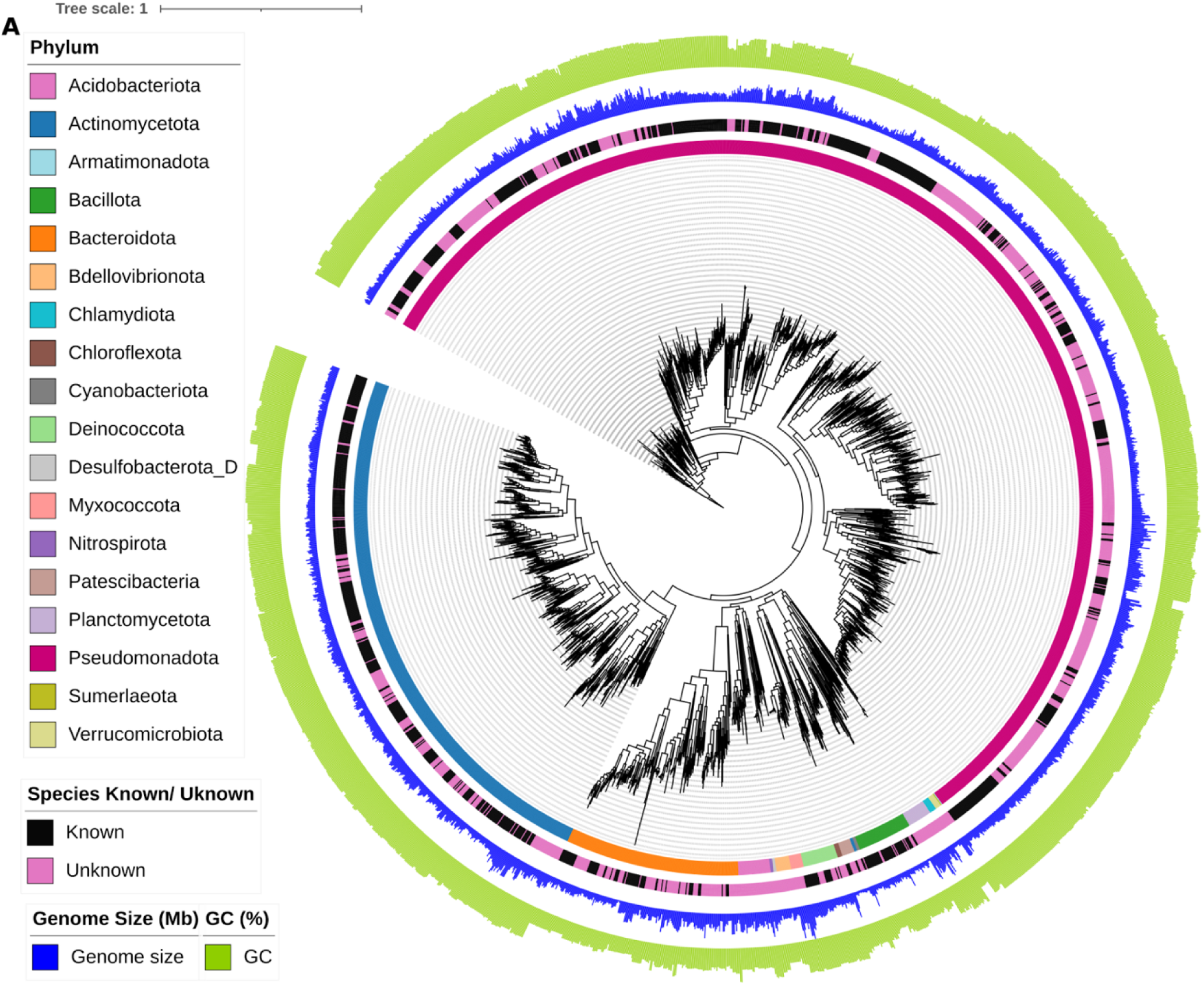
Phylogenetic tree of reconstructed metagenome-assembled genomes (MAGs), illustrating their taxonomic diversity and genomic characteristics. The tree represents the genome-resolved phylogeny, with branches corresponding to individual MAGs. The first outer ring denotes taxonomic classification at the phylum level, highlighting the dominance of major bacterial lineages including Pseudomonadota, Actinomycetota, Bacillota, and Bacteroidota, alongside several less abundant phyla. The second ring indicates species-level novelty, distinguishing MAGs assigned to known species from those lacking species-level classification, revealing a substantial fraction of previously uncharacterized genomic diversity within the indoor microbiome. The outer rings represent genome-associated features, including genome size (Mb) and GC content (%), demonstrating broad variability across phylogenetic groups. Notably, genome size and GC content exhibit lineage-specific patterns, reflecting underlying evolutionary and ecological adaptations.

### Strain-level diversity in prevalent MAGs has no house-wise stratification

The prevalent MAG subset (n=66) included species with at least 5 MAGs recovered across the three location group categories: living area, bathroom, and kitchen (**Supplementary Fig. 2**). While several species were confined to a single location group, the most prevalent taxa were detected across multiple indoor compartments. To assess whether these broadly distributed species were also structured at the strain level, we next analyzed phylogenetic and ANI patterns for the most prevalent MAGs. Among these, *Paracoccus marcusii* (n=56), *Ottowia* sp. 018060485 (n=53), and *Kocuria palustris* (n=44) were the most prevalent species in the dataset, with distributions across multiple locations rather than being restricted to a single house location (**Supplementary Fig. 2, Supplementary Fig. 3, Supplementary Fig. 4**). To assess whether these prevalent MAGs also showed strain-specific clustering with house locations, phylogenetic trees were generated for the three species with the most MAGs recovered. These trees were annotated with ANI matrices and metadata for location group and house number. For *P. marcusii*, the ANI matrix showed mostly lower pairwise ANI values (**Supplementary Table 3**), with the recovered MAGs visible within it, indicating some strain-level divergence among the genomes. The same pattern was observed for Ottowia sp. 018060485, which also showed more variable ANI values across comparisons (**Supplementary Fig. 3**, **Supplementary Table 2**). In contrast, *K. palustris* showed a more tightly clustered ANI pattern (**Supplementary Table 4**) with generally higher pairwise similarity among its MAGs, suggesting lower strain-level diversity within the house microbiome (**Supplementary Fig. 4**). Across all three species, the annotated trees show MAGs recovered from multiple houses and location groups, and the pattern does not show an obvious segregation by samples from individual house or location group at the strain level. However, a partial location-associated clustering was evident for the *K. palustris* MAGs, where MAGs from the living area were grouped together, and another small cluster was observed for the MAGs corresponding to the kitchen area (**Supplementary Fig. 5**). Similarly, *P. marcusii* also has two such small clusters in the kitchen area (**Supplementary Fig. 3**). Despite these location-linked groupings, no house-specific clustering of the MAGs was observed.

### Metabolic pathways in the house microbiome show spatial distribution along with a conserved core

The metabolic profile of the house MAGs was dominated by a conserved core of carbon and redox functions, with selective shifts across location groups (**Fig. 3e**). Differences in functional prevalence were assessed using Fisher’s exact test with Benjamini–Hochberg correction at an adjusted α of 0.05 for pair-wise testing across the location-groups. In the carbon cycle, organic carbon oxidation and fermentation were highly prevalent across all location groups, representing the core pathways (**Supplementary Table 6**). In contrast, acetate oxidation and carbon fixation were significantly more abundant in the bathroom (B) and kitchen (K) than in the living area (L) (FDR-adjusted p-value p < 0.001 for all). Carbon fixation potential was detected in 172 MAGs, spanning four pathways: the CBB (carbon benson-bassham) cycle with RuBisCO form I (138 MAGs), the 3-hydroxypropionate cycle (28 MAGs), reverse TCA cycle (5 MAGs), and CBB cycle with RuBisCO form II (1 MAG); at the genus level, these pathways were mainly represented by *Paracoccus*, *Amaricoccus*, *Erythrobacter*, *Aquincola*, *Mycobacterium*, *Aquabacterium_A*, *Sandaracinobacteroides*, *Blastomonas*, and *Nitrospira_A* (**Supplementary Table 7**). Methanotrophs were observed at lower abundances compared to the other categories, but were enriched in the bathroom region compared to the living area. In contrast, hydrogen oxidation and hydrogen generation were rare across all groups and did not differ significantly.

**Fig. 3.**
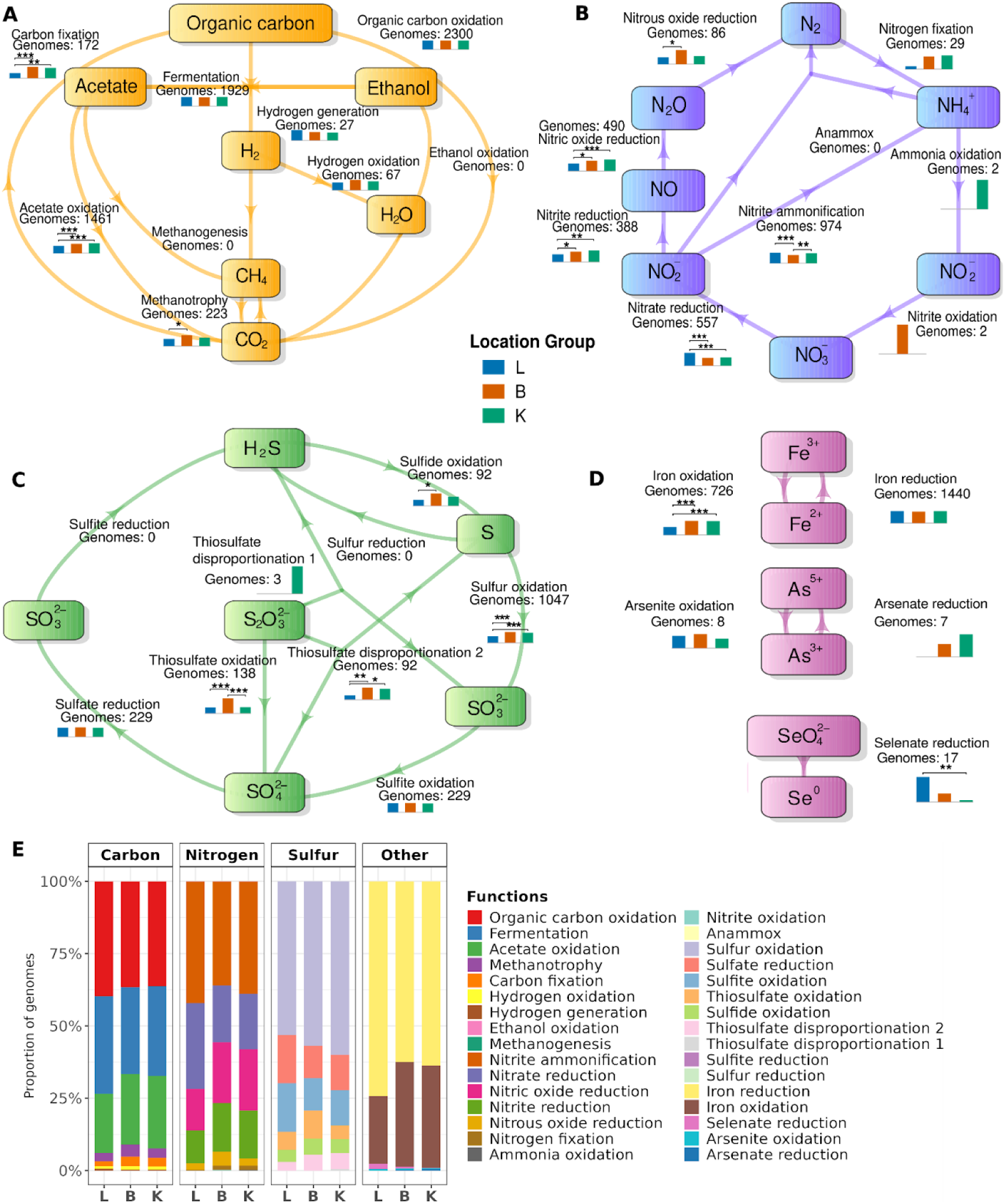
Functional potential of recovered MAGs reveals compartment-specific partitioning of carbon, nitrogen, sulfur, and other elemental cycling pathways. Schematic representation of key metabolic processes encoded by metagenome-assembled genomes (MAGs) recovered from the three location groups (L, B, and K). Carbon-related pathways include organic carbon oxidation, acetate oxidation, fermentation, methanotrophy, carbon fixation, hydrogen oxidation, hydrogen generation, ethanol oxidation, and methanogenesis. Nitrogen cycling encompasses nitrite/nitrate transformations, nitrogen fixation, ammonia oxidation, and anammox-related processes. Sulfur cycling includes sulfate, sulfite, thiosulfate, sulfide, and sulfur oxidation/reduction and disproportionation pathways. Additional functional capacities include redox transformations of iron, selenium, and arsenic. The stacked bar plot shows the proportions of genomes in each location group that encode the corresponding functional categories, highlighting differences in metabolic potential across indoor niches. Collectively, these results indicate that house-associated MAGs are functionally diverse and display location-dependent enrichment of pathways involved in elemental cycling and resource acquisition.

Nitrogen metabolism was inclined toward reductive transformations in the house microbiome. Nitrite ammonification was abundant in all groups, and nitrate reduction was enriched in L relative to B and K (190/570 versus 203/991 and 164/743; FDR-adjusted p = 9.38 × 10-8 and 9.07 × 10-6 for L vs B and L vs K, respectively). Nitrite reduction and nitric oxide reduction followed the same direction, with lower representation in L than in B and K. Nitrous oxide reduction was less abundant overall, whereas nitrogen fixation, ammonia oxidation, and nitrite oxidation were rare and insignificant. Sulfur and iron metabolism showed a similar spatial bias, with sulfur and iron oxidation more prevalent in the bathroom and kitchen. Sulfite oxidation, sulfate reduction, and iron reduction were broadly distributed across all the location groups.

### The living area has a distinct functional profile, whereas the bathroom and the kitchen are broadly similar

In the differential function enrichment analysis, the living area (L) showed the strongest separation from the other two location groups, whereas the bathroom (B) and kitchen (K) areas were more similar to each other (**Fig. 4a-c**). This was also evident in the number of functions that were significantly different between the groups after FDR correction: 36 in B versus L, 34 in K versus L, and only 6 in B versus K. The functions that distinguished L from B and K include chlorite reduction, nitrate reduction, lactate utilization, acetogenesis, pyruvate ⇌ acetyl-CoA + formate, Complex V (ATP synthase: V/A-type H+/Na+-transporting ATPase), aminotransferase class I and II, 4-aminobutyrate aminotransferase and related aminotransferases, selenate reduction, and perchlorate reduction. In contrast, B and K shared a common functional profile, both being enriched for cytochrome c oxidase, cbb3-type, Complex I (NADH-quinone oxidoreductase), Complex III (cytochrome c reductase), F-type ATP synthase, acyl-CoA dehydrogenase, cellulose degrading, chitin degrading, sulfur oxidation, and methanol oxidation when compared with L. The direct B vs K comparison showed only a small functional split: B was enriched for aerobic CO oxidation and nitrile hydratase, whereas K was enriched for cytochrome (quinone) oxidase, *bo* type, acetate => acetaldehyde, nitrite reduction to ammonia, and pyruvate ⇌ acetyl-CoA + formate. These results point to location-associated differences in the metabolic potential, with the strongest distinction observed between L and the other two location groups.

**Fig. 4.**
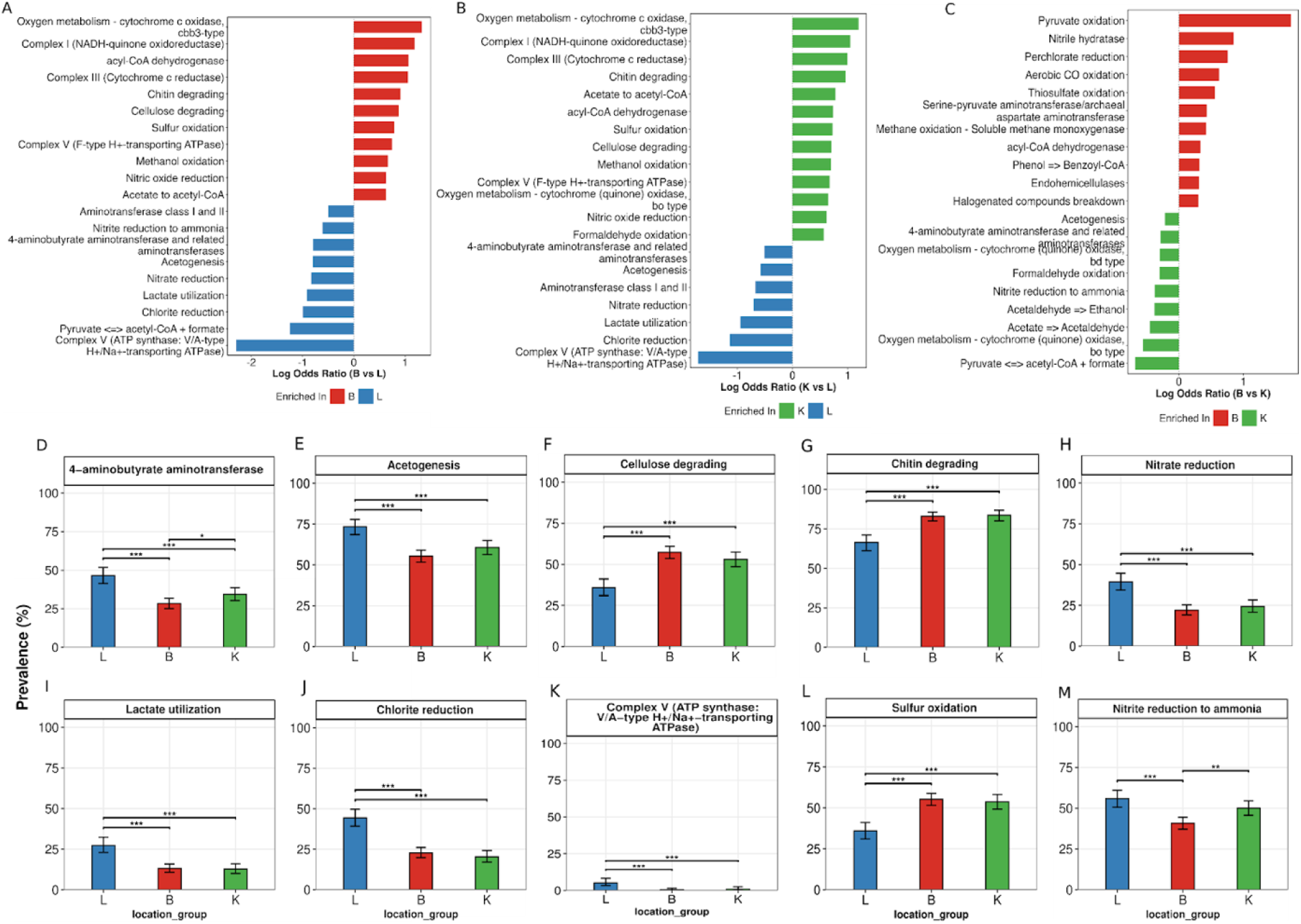
Location-dependent enrichment of metabolic traits across house microbiome MAGs. (a–c) Pairwise differential functional enrichment between bathroom (B), kitchen (K), and living area (L), shown as log odds ratios derived from 2 × 2 presence–absence tables. Positive and negative values indicate enrichment in the first- and second-named groups, respectively. Statistical significance was assessed using two-sided Fisher’s exact tests, and p-values were adjusted within each pairwise comparison using the Benjamini–Hochberg false discovery rate (FDR). (d–m) Prevalence of the same functions across B, K, and L, shown as the percentage of MAGs carrying each function, with 95% confidence intervals. Pairwise prevalence differences were tested using Fisher’s exact tests on the same presence–absence matrix, and p-values were corrected using the Benjamini–Hochberg FDR procedure. Brackets/asterisks denote FDR-adjusted p < 0.05.

Further, the prevalence of differentially enriched functions across the three location groups was tested (**Fig. 4d-m**). The prevalence differences were common and largely consistent with the enrichment patterns. Interestingly, among the 45 significantly enriched functions, 44 showed at least one significant prevalence across the location groups, indicating that the differential-enrichment signal was broadly mirrored by the distribution of functions across genomes rather than being driven by a small number of outlier MAGs.

We then explored the functional potential of the prevalent species MAGs relative to their reference genomes. Analysis revealed that the *K. palustris* showed the clearest species- specific functional differences compared to the reference genomes: chitin degradation was detected in 18/44 MAGs but present in all 10/10 reference genomes, cbb3-type cytochrome c oxidase was present in only 4/44 MAGs despite being present in all references, perchlorate reduction was detected in 17/44 MAGs versus 10/10 references, and ethanol-fermentation-associated function was present in 22/44 MAGs versus 10/10 references (**Supplementary Fig. 6**). In contrast, *P. marcusii* and *Ottowia* sp. 018060485 did not show loss of any specific functions across their MAG sets compared with the reference genomes (**Supplementary Fig. 7, Supplementary Fig. 8**). In the preliminary read-based analysis of the house microbiome, members of extremophilic microbial taxa Deinococcus and Thermus were identified(Awasthi et al., 2025). Here, we recovered 36 MAGs from the Deinococcus-Thermus phylum, representing 7 different species (*Deinococcus wulumuqiensis*, *D. reticulitermitis*, *D. fonticola*, *D. ficus*, *D. grandis*, *D. budaensis,* and *Deinococcus* sp. NW-56). Within this clade, specifically, urea utilization was restricted to a subset of MAGs, including A43_bin.27, A40_bin.19, A73_bin.18, A89_bin.24, A37_bin.15, A18_bin.15, A74_bin.11, A36_bin.30, A28_bin.3, A19_bin.7, A64_bin.10 and A9_bin.15 (Fig. S9). The remaining Deinococcus-Thermus MAGs lacked this function, along with all the reference genomes.

### Potential pathogens and resistome in the house microbiome

Using the CDC list of pathogenic species^18^, a total of 52 potential and opportunistic pathogenic species-level MAGs were identified across the house locations, but their distribution was clearly non-uniform (**Supplementary Fig. 10**). Some species were recovered across multiple locations, whereas others were restricted to a single location. Species such as *Escherichia coli*, *Acinetobacter junii*, *Achromobacter xylosoxidans*, *Mycobacterium smegmatis*, *Staphylococcus epidermidis*, *Staphylococcus saprophyticus*, *Streptococcus salivarius*, and *Stenotrophomonas maltophilia* contributed to the detected possible pathogenic pool (**Supplementary Fig. 4**). The phylogenetic analysis extended this pattern to the broader pathogenic MAG collection (**Supplementary Fig. 9**), which was distributed across three major phyla: Proteobacteria, Firmicutes, and Actinobacteriota. The tree shows that pathogenic genomes were phylogenetically diverse rather than concentrated in one narrow lineage, spanning genera including *Escherichia*, *Acinetobacter*, *Enterobacter*, *Pseudomonas*, *Stenotrophomonas*, *Achromobacter*, *Staphylococcus*, *Streptococcus*, *Enterococcus*, *Mycobacterium*, *Kocuria*, *Corynebacterium,* and *Microbacterium* (**Fig. 5**). The bar chart annotation in the phylogenetic tree shows uneven ARG burden. Only a subset of pathogenic MAGs carried a relatively high number of ARGs, whereas many others carried one or a few ARGs. This indicates that resistance potential within the pathogenic MAG set was lineage-specific and strongly skewed toward a limited number of taxa. Taxa with high ARG counts include *E. coli*, *Pseudomonas aeruginosa*, *Leclercia adecarboxylata*, and emerging drug-resistant pathogens like *Stenotrophomonas maltophilia*.

**Fig. 5.**
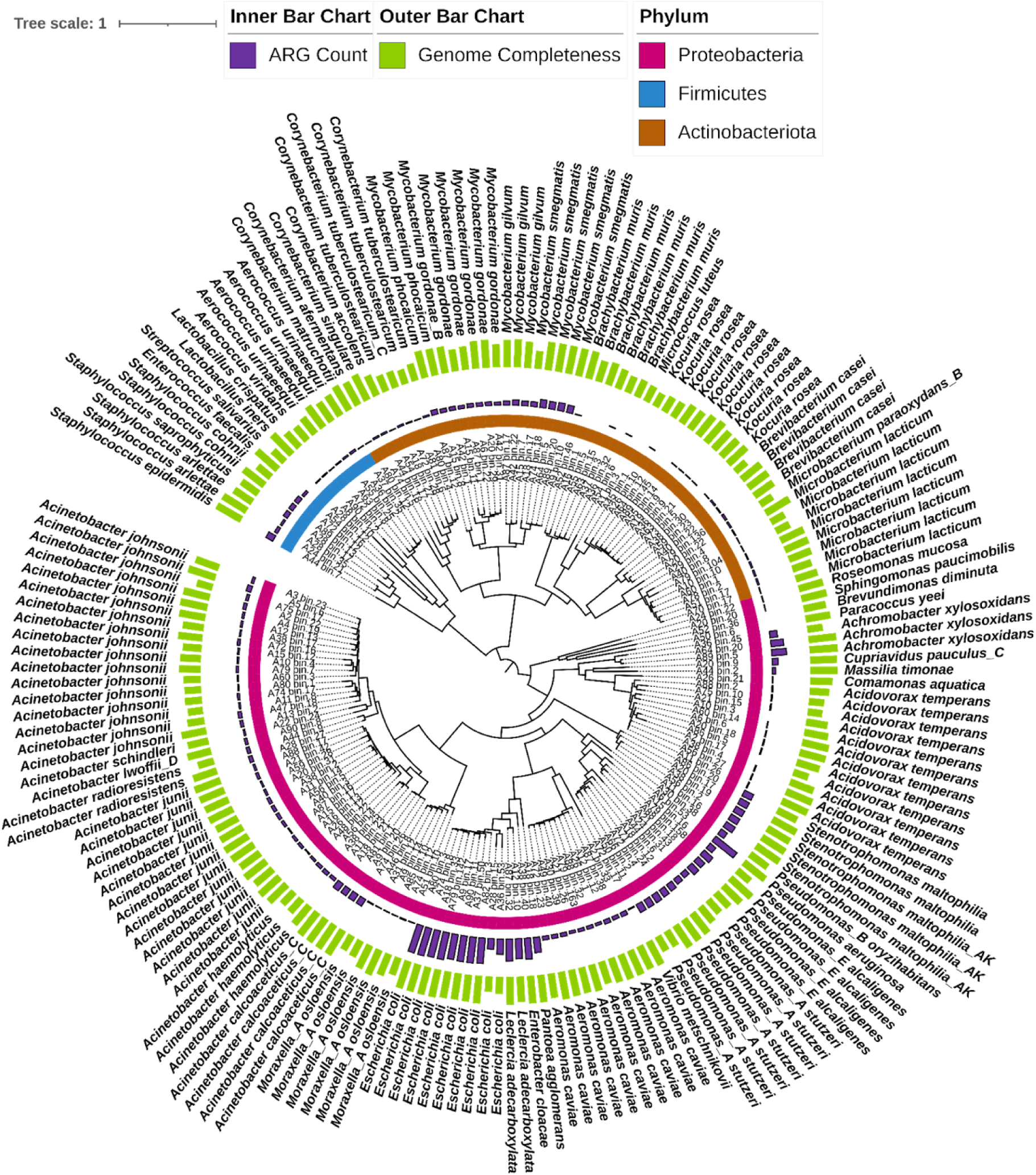
Phylogenetic distribution and genomic features of pathogenic MAGs. Circular phylogenetic tree of pathogen-associated metagenome-assembled genomes (MAGs) recovered from the house microbiome. Branches represent individual MAGs and reference genomes, colored by phylum as indicated. The inner bar plot shows the number of antimicrobial resistance genes (ARG count) detected in each genome, and the outer bar plot shows genome completeness. The tree highlights the phylogenetic placement of pathogenic MAGs across major bacterial phyla, together with genome quality and ARG burden, supporting genome-resolved identification of clinically relevant lineages in indoor environments.

**Fig. 6.**
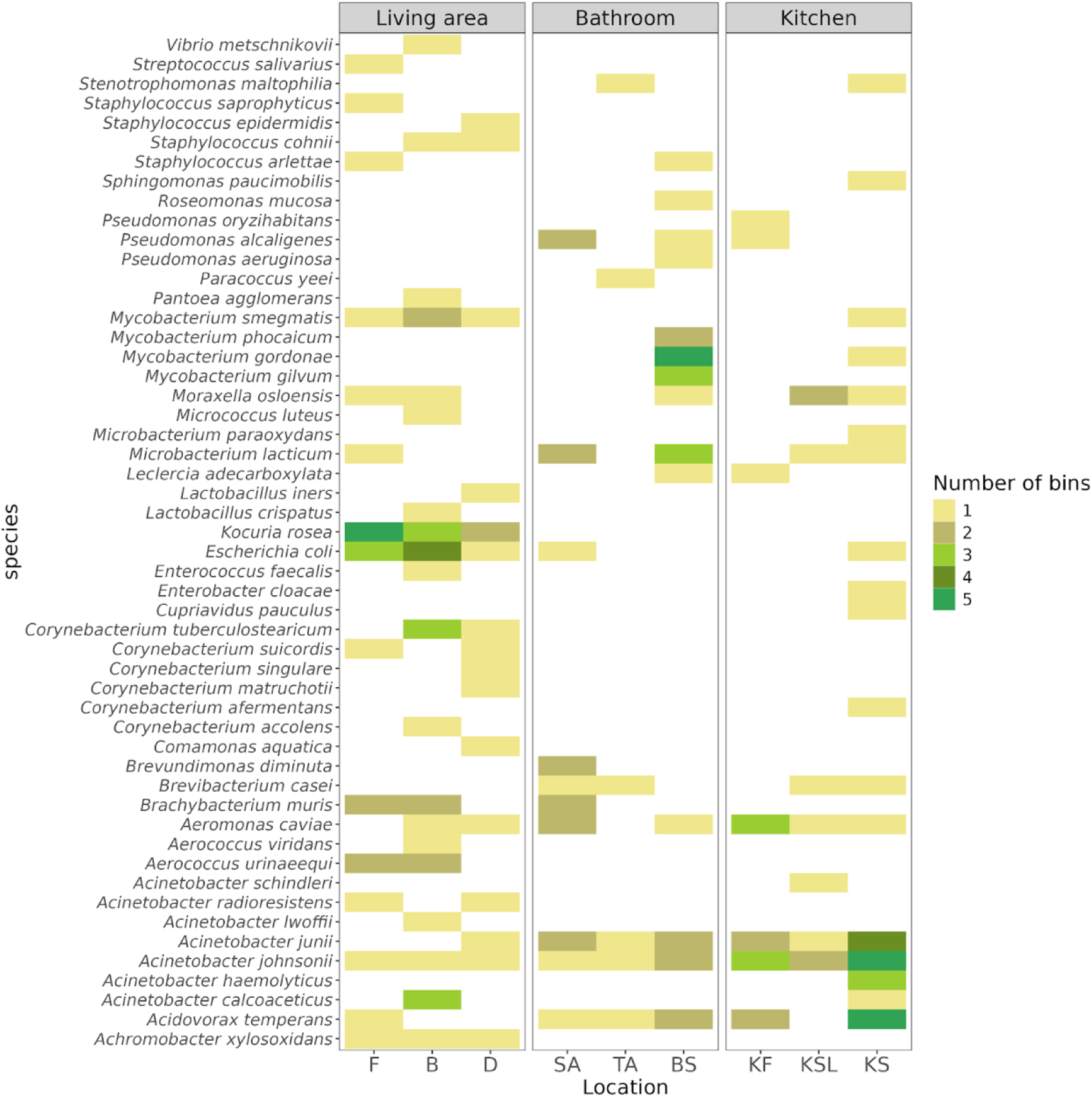
Distribution of pathogenic MAGs across house locations based on the CDC pathogen list. Heatmap showing the occurrence of pathogenic species-level metagenome-assembled genomes (MAGs) across living area, bathroom, and kitchen sampling locations. Rows represent pathogenic species included in the CDC-based pathogen list, and columns correspond to individual house locations within each compartment. Color intensity indicates the number of MAGs recovered per species at each location.

A total of 1080 ARG hits were detected in 440 house MAGs, representing 196 unique ARGs. The overall resistome was dominated by multidrug resistance, with 488 hits, followed by bacitracin, beta-lactam, and aminoglycoside, with 145, 139, and 79 ARG hits, respectively (**Supplementary Fig. 11a**). At the mechanistic level, efflux-associated resistance was the predominant mode, with 537 hits, indicating that the resistome is largely driven by broad-spectrum resistance mechanisms (**Supplementary Fig. 11b**). A total of 64 mobile genetic elements (MGEs) were identified on contigs carrying ARGs across 45 MAGs (**Supplementary Table 13**). Of these, 30 MGE-associated ARGs were clinically relevant, and 26 ARG-MGE pairs had less than 10 kbp distance between them (**Supplementary Fig. 12**). Insertion sequences accounted for most ARG-MGE associations (49/64), followed by miniature inverted-repeat transposable elements (MITES) (11/64), composite transposons (3/64), and single integrative conjugative elements. Plasmid association was rare and observed only for a single MAG, A16_bin.15, which was identified as *Psychrobacter faecalis* by GTDBtk. On the contig k141_460342 of the A16_bin.15, the aminoglycoside (*AAC (3’)-II*) resistance was linked to the insertion sequence ISAba14, and the same contig was also predicted as a plasmid (**Supplementary Table 17**).

The ARG–host network, detected among the MAGs, showed a hub-centered rather than diffuse architecture (**Fig. 7**). *E. coli* formed the largest central module in the network, but the broader structure was driven by a small number of highly connected resistance determinants, especially the efflux- and transport-associated genes *acrAFE*, *tol*C, *mdt*FHMKL, *emr*BDY, *mex*DFH, *opm*H, together with *sul*2, *bac*A, *arn*A, *ept*A and *ugd*. Several organism-specific associations were also found. The species, such as *Staphylococcus epidermidis,* carried *aad*A5 and *mph*(C); *Rothia* MAGs were linked to *fus*B and TEM; *Flavobacterium* carried *lnu*(A); and *Acinetobacter* MAGs were linked to ANT-and APH-type genes. *Achromobacter xylosoxidans* and *Pseudomonas* MAGs also contributed to efflux-associated resistance genes. It is to be noted that clinically relevant ARGs were not confined just to the putative canonical pathogens, but they were also present in MAGs assigned to non-classically pathogenic or environmental taxa such as *Sphingomonas*, *Rothia*, *Flavobacterium*, *Pseudoxanthomonas*, *Mesorhizobium,* and related lineages, indicating that the house microbiome functions as a broader reservoir of clinically important resistance genes.

**Fig. 7.**
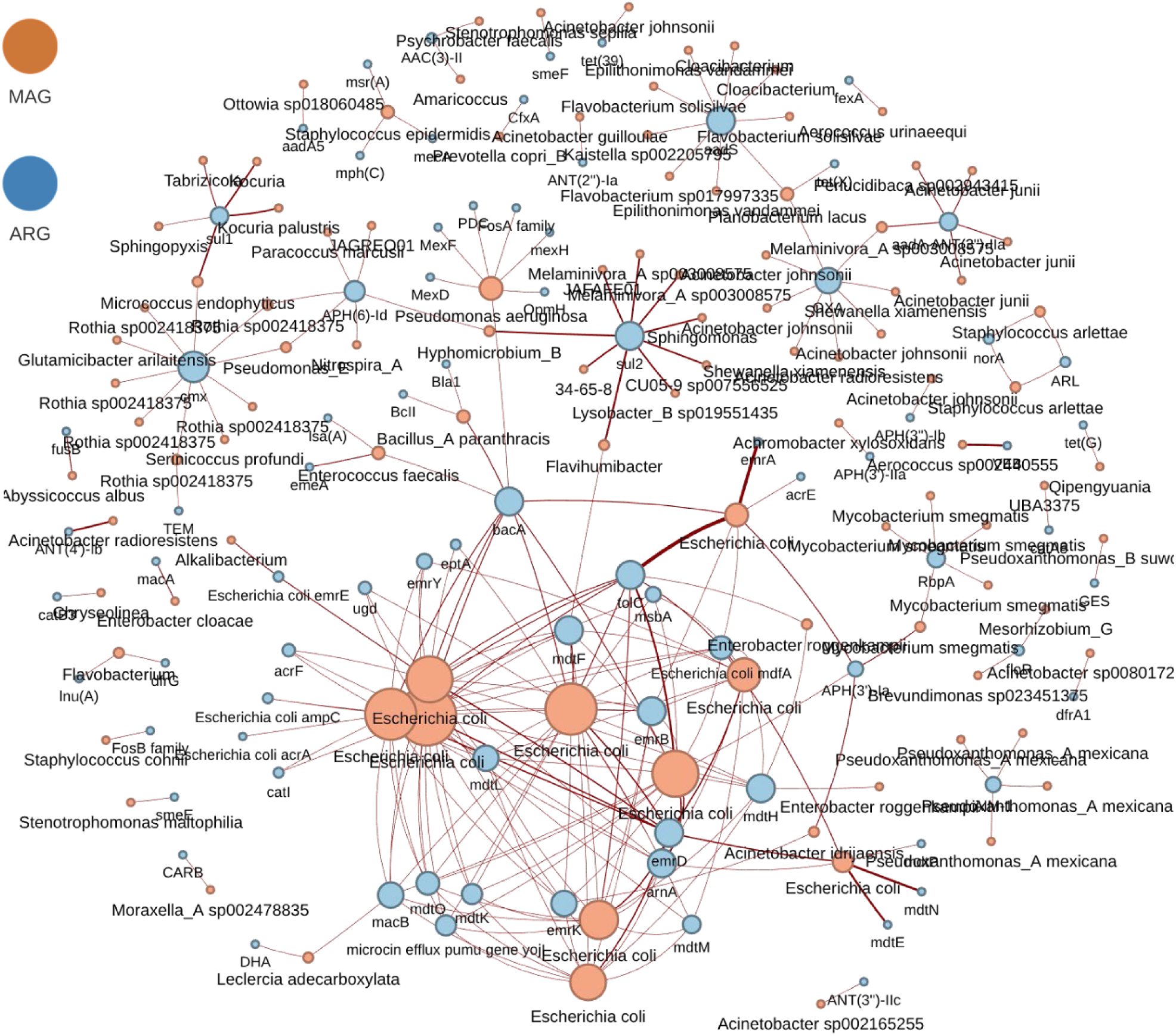
Network of MAG-ARG associations highlights potential genomic hosts of clinically relevant resistance determinants. Network visualization of metagenome-assembled genomes (MAGs) and antimicrobial resistance genes (ARGs) recovered from the house microbiome. Orange nodes represent MAGs and blue nodes represent ARGs; node size reflects connectivity, with larger nodes indicating genomes or genes linked to a greater number of associations. Red edges denote MAG ARG links, suggesting potential carriage of resistance genes by specific genomes. The network reveals a highly connected central module dominated by *Escherichia* and other taxa, alongside multiple smaller subnetworks, indicating that ARGs are unevenly distributed across microbial hosts and that a subset of genomes may act as key reservoirs of antimicrobial resistance within indoor environments.

### Phylogenomic analysis of the *E. coli* MAGs reveals affiliation with fecal commensal and pathotype strains

Of the 10 *E. coli* MAGs recovered from the house microbiome, 8 met the MIMAG standards for near-complete bins and were used for subsequent analyses. These MAGs were distributed across multiple household areas, including the drawing room, foyer, bedroom, kitchen sink, and shower area (**Supplementary Fig. 10**). To infer the likely origin of the recovered *Escherichia coli* MAGs, we performed a phylogenomic analysis using a curated reference set of pathogenic, commensal, and environmental strains^11^. Whole-genome phylogenetic analysis of *E. coli* MAGs recovered from the house microbiome revealed a dichotomy in the evolutionary origins and pathogenic potential of the house *E. coli* MAGs (**Fig. 8**). The resulting tree placed the eight MAGs within the broader *E. coli* phylogeny, rather than as a distinct house-specific lineage. Importantly, MAGs originating from different locations within the houses appeared to be intermixed within the tree and mostly belonged to one of the two major phylogroups of *E. coli*: group A and group B1. Five MAGs (A18_bin.33, A74_bin.8, A9_bin.17, A37_bin.50, and A99_bin.11) had close genetic relationships with the reference InPEC strains, while three (A79_bin.142, A82_bin.3, and A80_bin.3). had similarities with commensal fecal isolates The closely related commensal strains include ECOR01 and CIP61.11 while the pathogenic strains include 101-1(InPEC), and 55989(In-PEC)^19^. None of the MAGs clustered with the environmental strains, negating their environmental origin and indicating a plausible fecal origin.

**Fig. 8.**
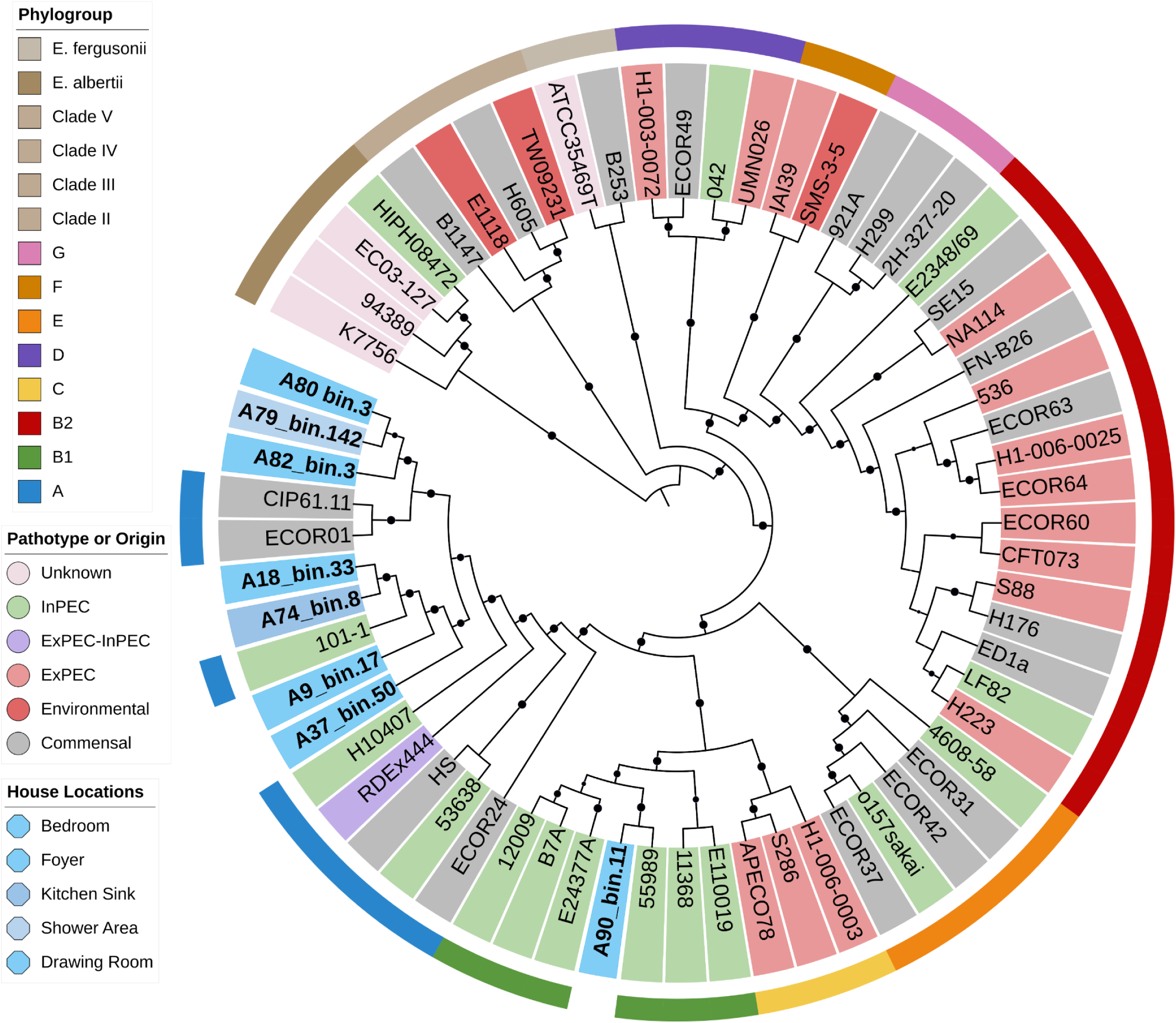
Phylogenetic placement and pathotype classification of *Escherichia coli* MAGs from the house microbiome. Circular phylogenetic tree showing the relationship of *E. coli* metagenome-assembled genomes (MAGs) recovered in this study with reference strains representing major phylogroups and *Escherichia* clades. Branches are colored according to phylogroup or species classification, including groups A, B1, B2, D, E, F, G, and cryptic clades, as well as *E. albertii* and *E. fergusonii*. Inner annotations indicate individual genomes, including MAGs and reference strains, while the outer rings denote pathotype or origin classification (including intestinal pathogenic *E. coli* (InPEC), extraintestinal pathogenic *E. coli* (ExPEC), commensal, environmental, and unknown categories) and sampling location within the house (drawing room, foyer area, bedroom, kitchen sink, and shower area). The size of the black circles indicates the bootstrap.

Further, ARGs and virulence factors were identified in the genomes to assess pathogenicity (**Fig. 9**). The ARG screening identified widespread carriage of resistance determinants, dominated by multidrug efflux systems. Core efflux and transport-associated genes such as *tolC, acrF, macB, emrY, mdfA, emrD, emrB, mdtO, mdtF, mdtL, msbA*, and the microcin efflux gene *yojI* were widespread across the MAGs. In addition, several clinically relevant resistance determinants were detected, including *amp*C (β-lactam resistance), *mph*(B) (macrolide resistance), *APH (3’)-Ia* (aminoglycoside resistance), and *catI* (chloramphenicol resistance)^20^. Envelope-modification genes linked with resistance to cationic antimicrobial peptides and polymyxin-like stress, including *arn*A, *ugd*, *ept*A were identified; in addition, *bac*A, an antimicrobial peptide resistance-associated transporter, was also detected^21–24^.

**Fig. 9.**
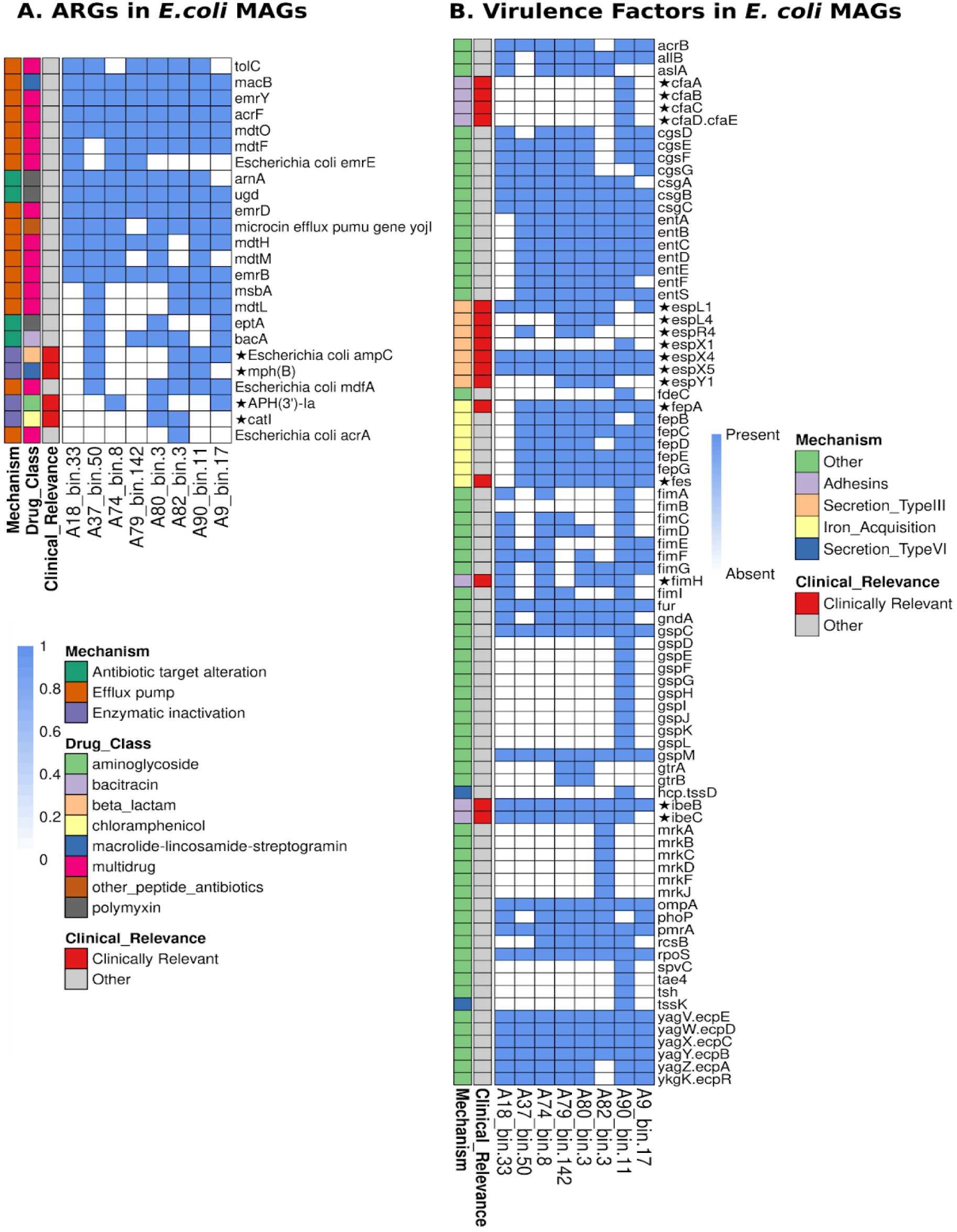
Distribution of antimicrobial resistance and virulence determinants in *Escherichia coli* MAGs. Heatmaps showing the presence and absence of antimicrobial resistance genes (ARGs) (left) and virulence-associated genes (right) across *E. coli* metagenome-assembled genomes (MAGs) recovered from house environments. Columns represent individual MAGs and rows correspond to genes grouped by functional categories. Blue indicates presence and white indicates absence. For ARGs, genes are classified by resistance mechanism (antibiotic target alteration, efflux pump, enzymatic inactivation) and associated drug classes, with clinically relevant genes highlighted. The profiles reveal widespread occurrence of multidrug efflux systems and resistance determinants across MAGs, alongside variability in specific antibiotic classes. For virulence-associated genes, functions are grouped into adhesins, secretion systems (Type III and Type VI), iron acquisition systems, and other factors. Several MAGs encode multiple virulence determinants, including adhesion factors, siderophore-associated iron-uptake systems, and secretion-related proteins, suggesting potential for host interaction and pathogenicity.

Analysis of virulence factors revealed a mixed virulence profile. The genes involved in colonization and persistence were widely distributed across the MAGs and included curli/fimbrial genes and enterobactin-based iron acquisition genes (*fepA, fepB, fepC, fepD, fepE, fepG,* and *fes*)^25–29^. The outer membrane and stress-responsive regulators (*ompA, phoP, pmrA, rcsB,* and *rpoS*), and the *E. coli* common pilus locus (*yagV/ecpE, yagW/ecpD, yagX/ecpC, yagY/ecpB, yagZ/ecpA,* and *ykgK/ecpR*) were also present^30–35^. The pathotype-associated virulence determinants were identified in a subset of MAGs and included colonization factor antigens (*cfaA, cfaB, cfaC, cfaD,* and *cfaE*), effector proteins (*espL, espR, espX,* and *espY*), and genes involved in invasion (*ibeB* and *ibeC*)^36–42^. VirulenceFinder, used as a complementary *E. coli*-specific virulence screen, recovered a narrower but more pathotype-focused set of determinants. Across the MAGs, canonical markers of colonization and persistence, such as fimH, csgA, hlyE, iss, ompT, shiA, terC, and yghJ, were detected, while A90_bin.11 carried the richest virulence profile, including *astA*, the complete *fae* locus, *fdeC, lpfA, tia,* and the *yehA-D* cluster. This supports A90_bin.11 as the strongest intestinal-pathotype-associated MAG in the dataset, whereas the remaining MAGs mainly showed adhesion, persistence, and stress-survival traits. Among the five *E. coli* MAGs that clustered with pathotype-associated reference strains in the phylogenetic tree, all five carried clinically relevant virulence factors, whereas four of the five also harbored clinically relevant ARGs. Specifically, A18_bin.33 contained pathotype-associated VFs, but no ARGs; A74_bin.8 carried *APH(3’)-Ia*; A9_bin.17 carried *amp*C, and *APH(3’)-Ia*; A37_bin.50 carried *amp*C and *mph*(B); and A90_bin.11 carried mph(B) and ampC, along with the richest virulence repertoire in this set. Importantly, the commensal cluster MAGs were not devoid of ARGs: A80_bin.3 contained *catI* and *APH(3’)-Ia*, while A82_bin.3 carried *catI, ampC,* and *mph(B)*. ARG-associated MGEs in the *E. coli* MAGs were dominated by insertion-sequence- and MITE-associated neighborhoods. The mobile element-linked ARGs included *APH(3’)-Ia* in close proximity to insertion sequences, typically within 10 kbp. The MAGs clustering with the pathogenic strains A9_bin.17 and A74_bin.8 carried *APH(3’)-Ia* in IS26-associated composite transposon contexts.

## Discussion

The present study is one of the largest genome-resolved house microbiome datasets reported yet. These micro-environments are structured microbial ecosystems varying with the locations of the houses, which were reported in our previous study^12^. Extending our previous work demonstrating dual environmental and human contributions to the house microbiome, this study delivers genome-resolved insights into strain-level diversity, metabolism, and clinically relevant features.

Recovery of 2304 MAGs, including 75 high-quality draft and 662 near-complete MAGs, indicates that a substantial portion of the house microbiome could be resolved at the genomic level. The highest number of MAGs was recovered from the bathroom location group, followed by the kitchen and living areas. The taxonomic distribution of the MAGs aligns with the read-level analyses from the pilot study of the New Delhi house microbiome. The genomic comparison using ANI vs. AF suggests that the recovered genomes are not phylogenetically or evolutionarily uniform. These represent phylogenetically divergent and low-abundance members of the indoor microbiome that escape species-level assignment, highlighting a substantial reservoir of previously uncharacterized household diversity. Their presence suggests that indoor microbial communities are not dominated solely by well-resolved core taxa, but also include dispersed, evolutionarily distinct lineages that may contribute specialized functional traits or transient ecological inputs^43–45^. This view is further supported by the taxonomic assignment, where nearly half of the MAGs remain unclassified at the species level.

The prevalent MAGs of the *P. marcusii* and *Ottowia* sp. 018060485 reveal substantial within-species diversity while remaining functionally similar to their reference genomes. This may suggest that the presence of these species is an input from the environmental sources with limited selective constraint^46,47^. *Paracoccus* species are often associated with healthy skin microbiota^48^, and their high prevalence may reflect metabolic versatility, including the potential for CBB RuBisCO form I–mediated carbon fixation, which could confer an advantage in nutrient-poor house environments. In contrast, *K. palustris* shows comparatively lower strain-level variation, as reflected by limited divergence in ANI values among its MAGs and by consistent functional differences with its reference genomes. This species occupies a broad ecological niche as the reference genomes were isolated from diverse habitats, including the International Space Station, cleanroom, soil rhizosphere, swimming pool water, human duodenal mucosa, and environmental sources^49–52^ (**Supplementary Table 2**). In particular, functions for the complex carbon degradation for chitin, perchlorate reduction, ethanol fermentation, and *cbb*3-type cytochrome c oxidase were less prevalent in MAGs. Interestingly, *cbb*3-type cytochrome c oxidase is a high-affinity terminal oxidase that supports respiration under low oxygen conditions and has been linked to fitness advantages in microaerobic and biofilm-associated lifestyles in bacteria^53,54^. However, other terminal oxidases, such as the Caa3 and bd types, were present across all MAGs and the reference genomes, indicating that respiration is maintained through alternative pathways^55^. The reduced prevalence of the *cbb*3-type oxidase may suggest the differences in oxygen availability in the house environment, where more stable oxygenated conditions could reduce the selection for high-affinity microaerobic respiration. Shifts in terminal c-oxidase usage have been linked to adaptation to oxygen gradients and ecological niches in bacteria^56^. This may imply a constrained population structure, potentially indicative of niche-specific adaptation to the house environment. Despite the strain level and functional differences among the prevalent MAGs (**Supplementary Fig. 2, Supplementary Fig. 3, Supplementary Fig. 4**), only limited clustering was observed based on location-group, not house-specific, as observed in the phylogeny. This suggests that the observed strain diversity is not clearly driven by houses, but some strains may have a preferential adaptation to the micro-environmental conditions at the locations of the houses. *Deinococcus*-related taxa have previously been reported from house microbiomes^57^, yet the ecological basis for their persistence in indoor environments remains unclear. Their recovery is consistent with the well-known stress tolerance of this phylum, including resistance to desiccation and other harsh conditions^58^ that are common on indoor surfaces due to low nutrient availability. Notably, urea-utilization genes were detected only in the house MAGs and were absent from all available reference genomes, suggesting a possible adaptation to occupant-derived urea from skin shed and sweat^59^ within the built environment.

The built environment, especially, is considered oligotrophic due to limited nutrient supply in most indoor spaces^60^. Reviews describe built environments as spaces that receive microbial inputs from outdoor sources and human substrates, but only a subset of these survive the dry, desiccation-prone, and low-resource conditions^1,5^. Consistent with the background, the house MAGs show a conserved carbon metabolic core rather than a broad range of carbon-cycling pathways. Organic carbon oxidation and fermentation were retained across all three location groups, indicating that carbon use is a common survival strategy in the house MAGs (**Fig. 3**). This is consistent with the idea that microbes in oligotrophic environments maintain a small subset of core metabolic functions rather than broad catabolic repertoires. In such nutrient-limited conditions, bacteria survive by slow growth, baseline maintenance, and streamlined functions, or by losing non-essential traits. Apart from that, carbon fixation in the house floor microbiome is a minor, lineage-specific trait rather than a dominant function. It is mainly driven by a few metabolically flexible microbes, primarily using the CBB cycle, with limited contributions from other pathways like 3HP (3hydroxypropionate)^61–63,60^ and the rTCA (reverse tricarboxylic acid) cycle. This suggests that CO₂ fixation here supports survival under low-nutrient conditions rather than functioning as a primary production system. Methanogenesis is completely absent in all the MAGs. Bathroom and kitchen samples were relatively enriched for acetate oxidation, carbon fixation, and methanotrophy, while the living area was relatively depleted in those functions. The absence of methanogenesis implies that acetate oxidation can be considered acetate consumption; rather than syntrophy^64,65^. In other words, both the bathroom and kitchen seemed to promote the processing of small organic molecules, while in the living space, bacteria focused primarily on survival using already deposited organic matter. This idea fits in well with the understanding of the indoors as oligotrophic environments where bacterial activity is largely dependent on moisture and surface conditions^60^.

The Nitrogen cycling processes do not show homogeneous distribution across the location groups but are spatially enriched. The enrichment observed for the nitrate reduction in living area might be associated with the nitrogen metabolism performed by skin commensals like *Staphylococci* since many of these bacteria have nitrate reductase genes, allowing them to metabolize nitrate as an electron acceptor under anaerobic growth conditions; moreover, the predicted nitrate reductase gene of *Epidermidibacterium keratini* is known; however, in *Corynebacterium*, it may vary according to the strain/species^66,67^. Nitrite ammonification was the most dominant step, enriched in the living area and kitchen relative to the bathroom, plausibly as a nitrogen conservation and recycling strategy^68^. Simultaneously, enrichment of nitrous oxide reduction occurred in the bathroom compared to the living room, whereas enrichment of nitrite reduction and nitric oxide reduction occurred in both the bathroom and kitchen compared to the living room. This can hence be viewed as a location-specific split between the nitrogen-retentive and denitrification-type processes. DNRA preserves available nitrogen, whereas denitrification is modular and usually incomplete, as many microbes participate only in part of the process, each with its own intermediates^69^. So, the living area is related to the nitrogen-conservative side, while the bathroom and kitchen areas lie on the reductive side of the nitrogen cycle.

Sulfur and iron show a similar spatial redox pattern. The bathroom and kitchen showed enrichment in sulfur oxidation, thiosulfate disproportionation, and sulfide oxidation reactions, but sulfite oxidation and sulfate reduction showed a similar distribution across the MAGs from all locations. Similarly, iron reduction was common, whereas iron oxidation was more prevalent in bathrooms and kitchens. All these indicate that B and K exhibit more opportunities for redox interfaces, suggesting greater potential to participate in reduced sulfur metabolism and the oxidation of Fe (II), which is not the case in the living area. This interpretation is consistent with recent work showing that microbial sulfide oxidation can be coupled to iron (III) oxide respiration, linking sulfur and iron cycling through shared redox chemistry rather than treating them as independent pathways^70^.

Previous house-microbiome studies show that indoor communities are location-dependent and that built environments can act as reservoirs for both potential pathogens and antimicrobial resistance determinants^71–73^. In our house MAG dataset, the fact that only a subset of taxa carry the largest ARG loads argues against a diffuse resistome and instead points to lineage-specific resistance enrichment within a limited number of hosts^57^ (**Fig. 5**). Taxa like *Escherichia*, *Leclercia*, *Stenotrophomonas*, *Pseudomonas*, *Mycobacterium*, *Achromobacter*, and *Acinetobacter* harbor a larger number of ARGs than other potential/opportunistic pathogen MAGs (**Fig. 5**).

One particularly important conclusion from these findings is that the ARGs associated with pathogenicity were not limited to putative pathogens. The wider presence of ARGs in MAGs belonging to taxa such as *Sphingomonas*, *Rothia*, *Flavobacterium*, *Pseudoxanthomonas*, and *Mesorhizobium* indicates a broader pool of environmental resistance genes that could be retained in non-pathogenic hosts. Studies on indoor dust have indicated that ARGs can be plasmid-borne, mobile, and detectable in living bacteria, suggesting that the built environment provides an opportunity for the persistence and exchange of resistance determinants beyond the classical clinical pathogen^71,74^. Although it need not imply that all ARG-carrying environmental MAGs are hazardous, it does indicate that the microbial composition of a house can serve as a potential source of resistance genes that could be transferred to pathogens under the right ecological and selective conditions^75^. Mobile element-linked ARGs were identified in the MAGs, but not plasmid-linked ARGs, which may be due to technical and biological factors. Plasmid reconstruction from metagenomic short reads is often difficult, as plasmids are highly diverse and repeat-rich, which frequently fragment during assemblyA^76–78^. In our dataset, ARG-mobility could be attributed to insertion sequences and other non-plasmid MGEs, consistent with studies reporting that IS elements and integrons play major roles in ARG mobilization and dissemination^79–81^.

The MAGs for *E. coli* recovered from the house microbiome are inferred to be possible fecal-lineage-associated strains belonging to commensals, as well as pathotype-associated groups, rather than house-specific environmental populations. *E. coli* is a vertebrate gut commensal and opportunistic pathogen, whose pathogenicity is organized into pathotypes. Its virulence depends more on accessory genome elements, such as virulence factors and ARGs^19^.

Phylogenomically, the house *E. coli* MAGs were dominated by phylogroup A, with one in B1. This is consistent with the current distribution of fecal *E. coli*, since phylogroups A and B1 are frequently found among commensals or intestinal pathogens, while most extra-intestinal pathogens belong to phylogroups B2 and D; however, phylogroups A and B1 do not rule out pathogenicity^19^. The MAGs are clustered into two groups based on phylogenomic diversity: pathogenic (n=5) and commensal (n=3). For the MAGs in the pathogenic cluster, the main concern arises from the combination of virulence and clinically relevant ARGs within their genomes. For example, A74_bin.8 has APH (3’)-Ia, A9_bin.17 has ampC, and APH (3’)-Ia, A37_bin.50 has ampC and mph(B), while A90_bin.11 has mph(B) and ampC, with the highest virulence load among all MAGs in this cluster. Additionally, A18_bin.33 should be noted for its lack of ARGs found in the other MAGs. Regarding the biological plausibility of the comparison, the selected pathogenic context is realistic, as both EAEC and ETEC are diarrheagenic pathotypes, with ETEC strain E24377A being a confirmed human pathogen and EAEC strain 101-1 representing a well-known outbreak-associated strain in EAEC research^38,82^. From the commensal cluster MAGs, A80_bin.3 and A82_bin.3 possess clinically significant antibiotic resistance genes; however, A80_bin.3 is particularly significant due to the fact that the APH (3’)-Ia gene is found in a contig surrounded by transposases from the IS6/IS15 family. Usually, antimicrobial resistance in the environment is acquired through the transfer of bacteria or genes between humans, animals, and man-made settings, while mobile genetic elements are major drivers of ARG acquisition and assembly into resistance islands^75,83^.

From the above, it can be inferred that the house *E. coli* MAGs represent two layers of risk: the pathogenic cluster, which already combines virulence and resistance, and the commensal cluster, which may act as a mobile ARG reservoir. The concern is not only the direct pathogenicity but also the risk of disseminating clinically relevant ARGs into the community through horizontal gene transfer from fecal *E. coli* lineages. Apart from the *E. coli* MAGs, the presence of clinically relevant ARGs in non-classic environmental microbe hosts further underscores public health concerns in the household microbiome. The residential surfaces should not be considered solely as contact surfaces but also as microbial environments where potential pathogenic organisms, opportunists, and ARGs may coexist and interact. These findings extend the relevance of the house microbiome beyond the built environment and into the One Health framework, where indoor microbial communities may influence human exposure to pathogens, antimicrobial resistance, and mobile genetic elements.

This paper offers the genomic perspective on the house microbiome and illustrates that residential floor surfaces do not merely represent passive collections of microbes, but rather are niche-based, metabolically driven communities. The identification of 2,304 MAGs from New Delhi residential buildings has revealed considerable unexplored diversity, functional differentiation within the living spaces, bathrooms, and kitchens, and functional associations between various taxa and specific strain- and species-level metabolic processes. Moreover, the identification of the resistome associated with mobile carriers underscores that indoor spaces serve as a habitat for potential ARG dissemination. Finally, the recovery of *E. coli* MAGs from potentially pathogenic and commensal lineages underscores the importance of the indoor microbiome for public health. This study also lays the foundation for the development of bio-informed indoor infrastructures.

## Methods

### Study design and Sample collection

This study analyzed 90 shotgun metagenomic swab samples collected from 10 urban houses in New Delhi, India, comprising five houses with cemented floors and five with tiled floors. The houses were distributed across different regions of New Delhi and were occupied by 2–4 residents each. Samples were collected from nine household locations, including the foyer, bedroom, drawing room, shower area, toilet area, bathroom sink, kitchen floor, kitchen slab, and kitchen sink, one day after routine cleaning and before the next cleaning cycle. The sequencing data used for metagenomic analysis and MAG recovery were obtained from BioProject PRJNA1053252 and comprised a total of 800.8 Gb of Illumina NovaSeq 6000, 2x150 paired-end data, with per-sample yields ranging from 2.68 to 79.059 Gb. Human reads were removed using Bowtie2, which aligned each sample to the hg37 human reference genome. The sample with the maximum number of human reads was A90, in which 57% reads were mapped to the Hg37 reference genome, and the sample that had the minimum host DNA content was A88, in which only 182 reads were aligned to Hg37. The sample yield ranged from 1.4 Gbp to 73.12 Gbp in the A17 and A79 samples, respectively.

### Assembly and Binning

Assembly of each individual sample was performed using MEGAHIT v1.2.9^84^ (min_contig_len=1000 bp). Quality assessment of the assemblies was performed using QUAST v5.2.0^85^ (average total assembly size: 18.94 Gbp, average GC content: 61.87%, and average N50 value: 4366.68 bases). Binning was performed for the individual assemblies using three different metagenomic binning software: MaxBin2^86^, metaBAT2^87^, and CONCOCT^88^; After obtaining three sets of bins for each sample from three tools as mentioned above, the best bins were selected by MetaWrap^89^(v 1.2.1) software package using its bin_refinement module (parameters; c for completeness 50% and -x for contamination 10%), two genomes were identified as same if they have minimum of 80% overlap. Completeness and contamination parameters were determined according to the MIMAG standards developed by the Genomic Standards Consortium (GSC)^17^. Dereplication was performed for all the bins coming from each sample separately using dRep^90^ v3.4.0 according to average nucleotide identity (parameters: -com (completeness%): 50, -nc (coverage threshold defining the fraction of overlap between the genomes): 0.3, -con (contamination%):10, -sa (secondary ANI threshold%): 99. MAGs were dereplicated on a per-sample basis rather than across the entire dataset to preserve strain-level diversity. This approach ensures that distinct strains of the same species recovered from different spatial locations are retained for higher-resolution downstream analysis. To identify species-level genome bins (SGBs) across the house MAG dataset, all dereplicated MAGs from individual samples were further clustered using dRep v3.4.0 with a secondary ANI threshold of 95% (-sa 0.95), primary ANI threshold of 90% (-pa 0.90), minimum completeness of 50%, maximum contamination of 10%, and overlap coverage threshold of 0.30 for genome comparison.

### Quality Assessment of the MAGs

Refined bins obtained from metawrap were dereplicated and called Medium Quality Bins (completeness and contamination 50% and 10%, respectively). High-quality draft genomes were evaluated by first filtering out MAGs with 90% completeness and 5% contamination, and then identifying tRNA and rRNA gene sequences in each MAG. tRNA was detected using tRNAscan-SE v 2.0.7^91^, and rRNA gene sequences were detected using barrnap v 0.9^92^. MAGs were evaluated as high-quality draft Genomes, each containing at least 18 unique tRNAs and 16S rRNA, 23S rRNA, and 5S rRNA (partial or complete gene sequences). All these parameters for evaluating high-quality draft genomes were selected based on the MIMAG standards developed by the Genomic Standards Consortium (GSC). Near-complete MAGs were defined as those that had 90% completeness and contamination less than or equal to 5%.

### Taxonomy and Phylogeny of the MAGs

For assigning the taxonomy, the GTDB-Tk v 2.1.1^93^ classification workflow was used with release214 as the database. It classifies each genome based on its placement in the GTDBtk reference tree, its relative evolutionary divergence, and/or average nucleotide identity (ANI) to reference genomes. PhyloPhlAn v 3.0.60^94^ was used for MAG phylogenetic analyses (parameters: --diversity high -fast). The phylogenetic tree presented in this study was visualized using ITOL^95^.

### Antimicrobial resistance genes, Mobile Genetic Elements, and Virulence identification

ARGs were identified in all MAGs using the SARG^96^ database workflow. First, the SARG reference FASTA headers were standardized so that the DIAMOND^97^ subject identifiers matched the SARG structure file. This was done by rewriting the original FASTA header tokens to the corresponding SARG.Seq.ID values, generating a cleaned reference FASTA and a header-mapping log. The cleaned SARG reference set was then used to build a DIAMOND protein database. Each MAG was queried against this database using DIAMOND blastx with tabular output format 6. The DIAMOND output was processed using a custom script to filter, rank, and annotate ARG hits. Subject lengths were extracted from the cleaned SARG FASTA to calculate alignment coverage. For each alignment, subject coverage was computed as: coverage (%) = (alignment length/subject length) × 100. Only hits with ≥ 90% amino acid identity and ≥ 100% subject coverage were retained. For each contig within each MAG, only the best hit based on the highest bitscore was kept. The retained hits were then joined to the SARG structure file to assign functional annotations, including gene name, resistance type, and mechanism group. Clinically relevant ARGs were identified as listed in the study^20^ Virulence factors were detected using the VFDB database with the Abricate^98^ and VirulenceFinder^99^ tools. VFDB recovered a broader virulence repertoire, whereas VirulenceFinder highlighted a smaller set of canonical *E. coli* virulence markers, indicating that the two tools were complementary rather than redundant. The ARG-host network plot was constructed using the visNetwork v2.1.4^100^.

#### Functional annotation of the MAGs

For the functional annotation of the MAGs, the METABOLIC_G v 4.0^101^m tool was used with default parameters, and a pathway was considered present only when 75% of the associated genes were detected in that pathway.

### Differential functional analysis of MAGs

Metagenome-assembled genomes (MAGs) were filtered with completeness ≥70%, yielding 1,598 MAGs (724 B, 509 K, and 365 L). Functional annotations were mapped to MAGs and converted into a binary presence–absence matrix, such that a function was considered present if detected at least once within a MAG. Redundant function hits per MAG were collapsed, and functions absent from the filtered dataset were excluded, resulting in 69 functions retained for analysis. Differential functional prevalence was assessed using pairwise comparisons between location groups (B vs L, K vs L, and B vs K). For each function, a 2 × 2 contingency table was constructed based on presence–absence counts across groups and tested using a two-sided Fisher’s exact test, which is appropriate for discrete prevalence data and widely applied in genomic enrichment analyses. Effect sizes were summarized as log odds ratios, with a Haldane continuity correction applied where required. To control for multiple hypothesis testing, P values were adjusted within each pairwise comparison using the Benjamini–Hochberg procedure, and reported as false discovery rate (FDR)-adjusted p values (q values). Functions with q < 0.05 were considered significantly differentially enriched.

To assess whether differentially enriched functions were broadly distributed across location groups, we conducted a prevalence analysis on the subset of functions that were already significant in the differential functional analysis. Function profiles were extracted from the cleaned function matrix and converted to binary presence–absence values at the MAG level, with numeric values >0 and categorical indicators such as “present” treated as presence. Only high-quality MAGs with completeness ≥70% and assigned to the three target location groups (B, K, and L) were retained for the analysis. For each function within each location group, prevalence was calculated as the percentage of MAGs carrying that function, along with 95% confidence intervals for the proportion estimated using prop.test. To test whether function prevalence differed between location groups, pairwise 2×2 contingency tables were built from the number of present and absent MAGs for each function, and Fisher’s exact test was applied to each pairwise comparison. Raw p-values were adjusted using the Benjamini–Hochberg method, and adjusted p-values <0.05 were considered significant. The workflow was implemented in R using Fisher’s test for contingency table inference and the p-adjust method (method = “BH”) for multiple testing correction.

## Declarations

During the preparation of the manuscript, the authors used AI-assisted technologies for language improvement, followed by careful review and editing by the authors, who take full responsibility for the accuracy and integrity of the publication.

## Data Availability

The shotgun metagenomic sequencing data and associated sample metadata used in this study are publicly available in the NCBI Sequence Read Archive (SRA) under BioProject accession PRJNA1053252. The metagenome-assembled genomes (MAGs) generated and analyzed in this study have been deposited in the Zenodo repository https://doi.org/10.0.20.161/zenodo.20048476.

## Author Contributions

S.A. contributed to devising the methodology, performing data analyses, interpretation, writing the original draft & editing the manuscript. R.S conceived the concept, acquired funding, supervised, performed interpretation and writing & editing of the manuscript.

## Acknowledgements

We thank the Director of CSIR-IGIB for his constant support.

## Ethical Declarations

Not applicable

