## Supplemental figures for "Genome-resolved house microbiome exhibits location-specific metabolic partitioning, and harbors hosts with clinically relevant antibiotic resistance genes"

### Supplementary Figures

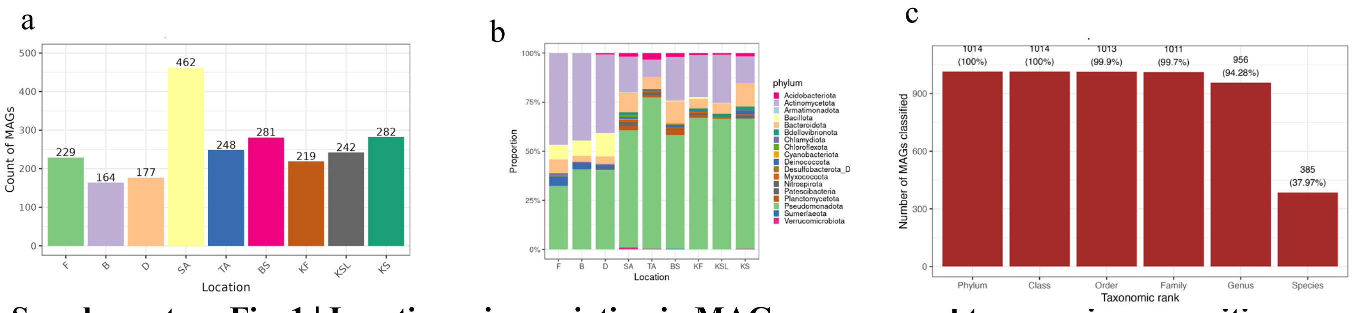


**Supplementary Fig. 1 | Location-wise variation in MAG recovery and taxonomic composition across house environments.**

(a) Number of metagenome-assembled genomes (MAGs) recovered from each sampling location, showing substantial variation in genome yield across sites, with SA contributing the highest number of reconstructed genomes, while other locations exhibit comparatively lower recovery. (b) Relative taxonomic composition of MAGs at the phylum level across locations. The community is consistently dominated by major bacterial phyla, including Pseudomonadota, Actinomycetota, and Bacillota, with location-specific differences in the relative abundance of less dominant lineages. Abbreviations for locations of the houses: F (Foyer), B (Bedroom), D (Drawing Room), SA (Shower Area), TA (Toilet Area), BS (Bathroom Sink), KF (Kitchen Floor), KSL (Kitchen Slab), KS (Kitchen Sink). (c) Taxonomic classification depth of dereplicated species-level genome bins (SGBs). Bar plot showing the number of SGBs classified at different taxonomic ranks based on GTDB-Tk annotation, from phylum to species level. Species-level classification was achieved for 385 SGBs, whereas 629 SGBs remained unclassified at the species level.

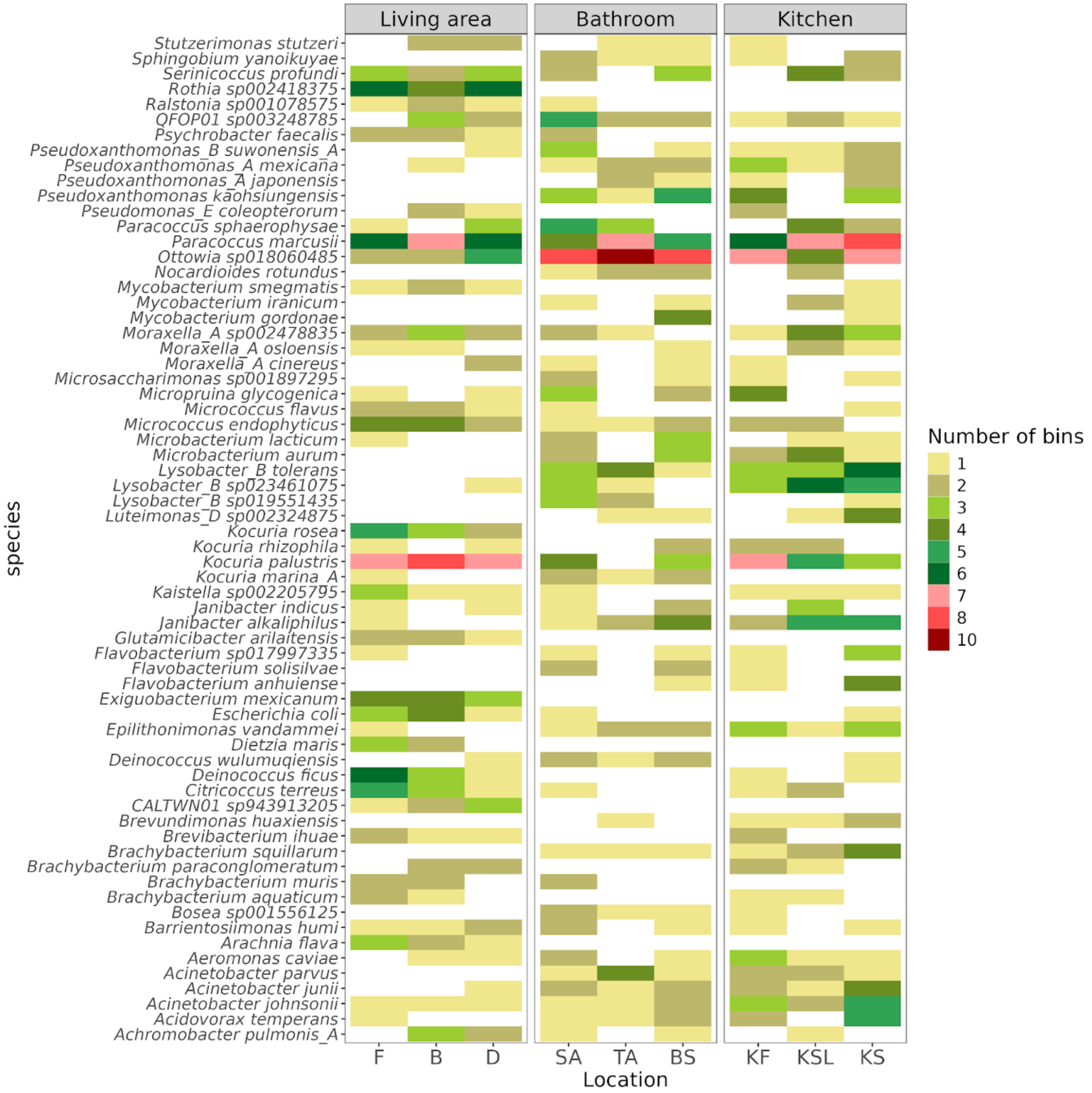


**Supplementary Fig. 2 | Species-level distribution of MAGs across house locations and compartments.**

Heatmap showing the distribution and abundance of species-level metagenome-assembled genomes (MAGs) across three major house compartments: living area, bathroom, and kitchen. Each row represents a species-level genome bin, and each column corresponds to a sampling location within each compartment. The color scale indicates variation in the number of bins recovered per species across the sampled locations.

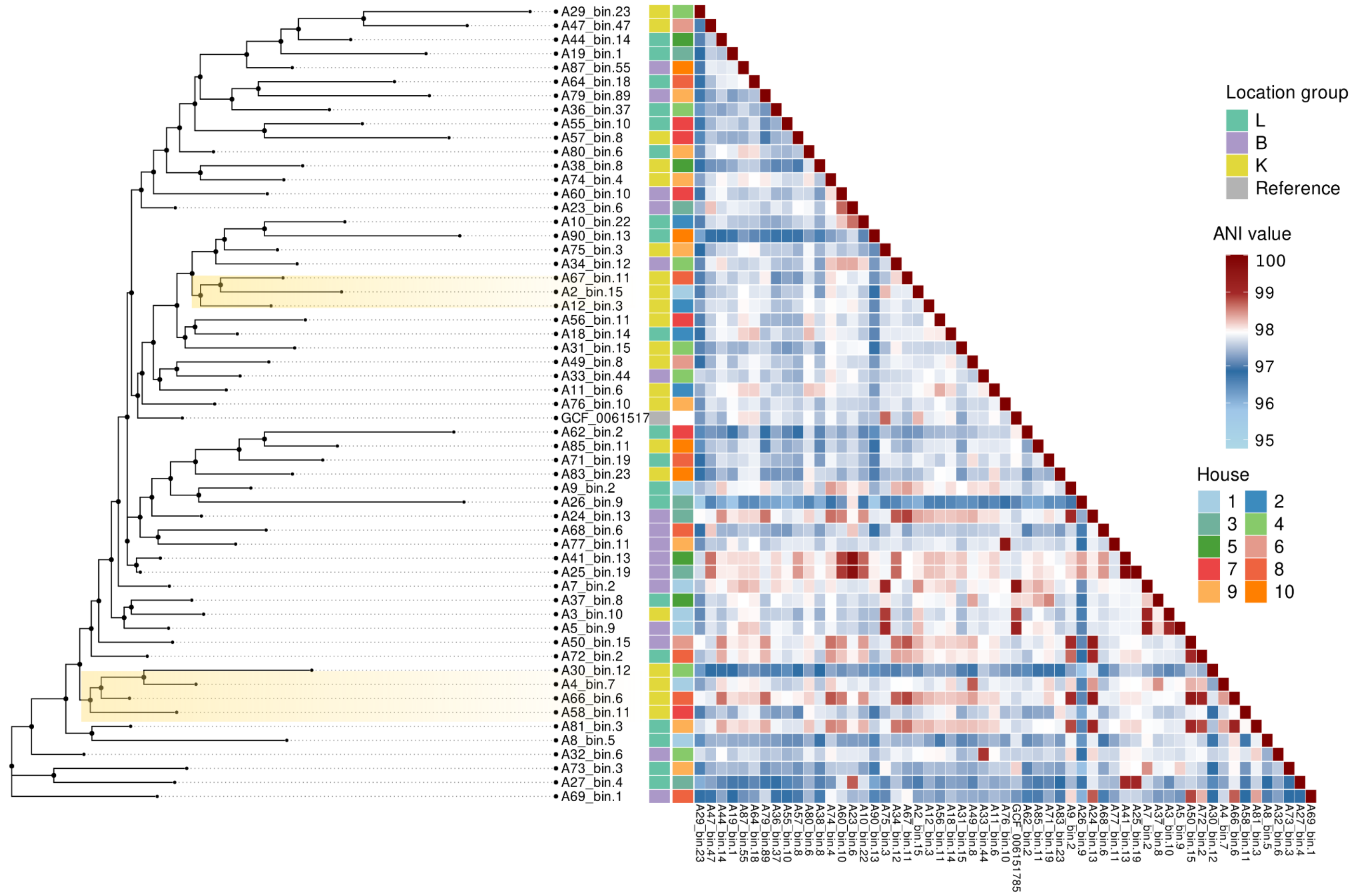


**Supplementary Fig. 3 | Strain-level variation among *Paracoccus marcusii* genomes recovered from the house microbiome.**

Pairwise average nucleotide identity (ANI) heatmap and hierarchical clustering of *Paracoccus marcusii* metagenome-assembled genomes (MAGs) and reference genomes. The dendrogram at left shows the phylogenetic relationships among genomes, while the triangular heatmap displays pairwise ANI values, revealing strain-level heterogeneity within the species. Genomes are annotated by location group (L, B, K) and house of origin, highlighting both within-location clustering and broader genomic relatedness across indoor sampling sites.

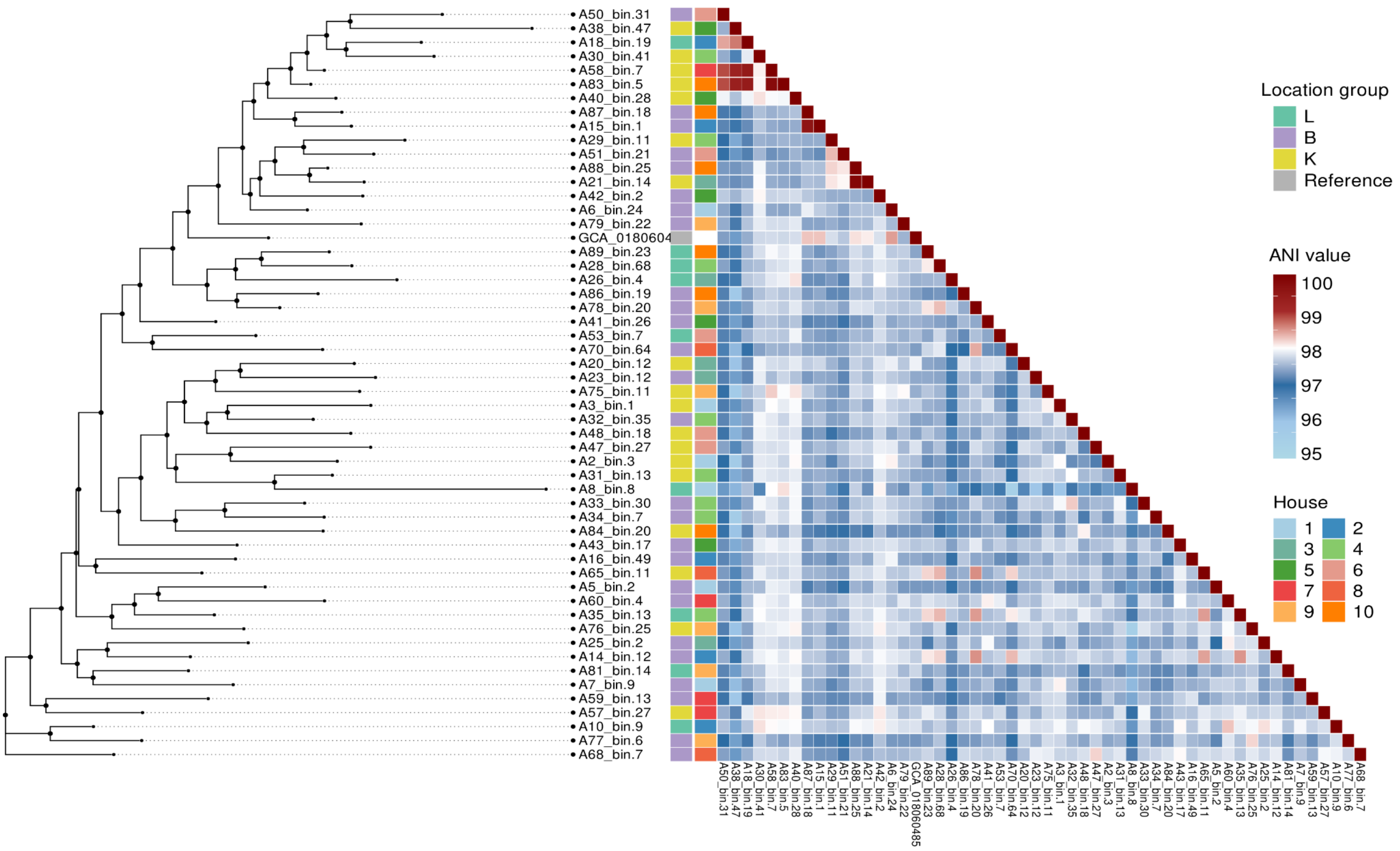


**Supplementary Fig. 4 | Strain-level variation among *Ottowia* sp. 018060485 genomes recovered from the house microbiome.**

Pairwise average nucleotide identity (ANI) heatmap and hierarchical clustering of *Ottowia sp* metagenome-assembled genomes (MAGs) and reference genomes. The dendrogram at left shows the phylogenetic relationships among genomes, while the triangular heatmap displays pairwise ANI values, revealing strain-level heterogeneity within the species. Genomes are annotated by location group (L, B, K) and house of origin, highlighting both within-location clustering and broader genomic relatedness across indoor sampling sites.

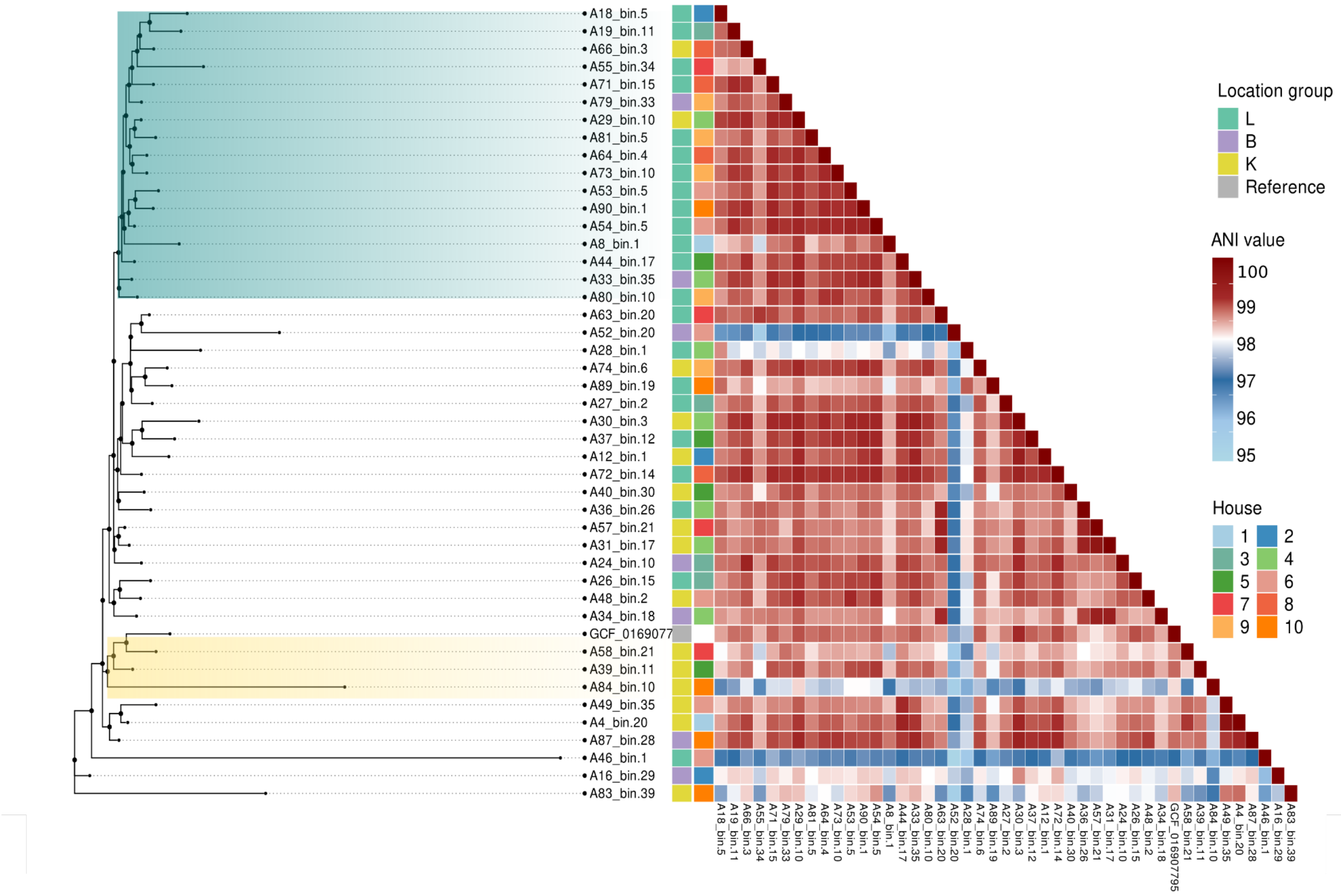


**Supplementary Fig. 5 | Strain-level variation among *Kocuria palustris* genomes recovered from the house microbiome.**

Pairwise average nucleotide identity (ANI) heatmap and hierarchical clustering of *K. palustris* metagenome-assembled genomes (MAGs) and reference genomes. The dendrogram at left shows the phylogenetic relationships among genomes, while the triangular heatmap displays pairwise ANI values, revealing strain-level heterogeneity within the species. Genomes are annotated by location group (L, B, K) and house of origin, highlighting both within-location clustering and broader genomic relatedness across indoor sampling sites

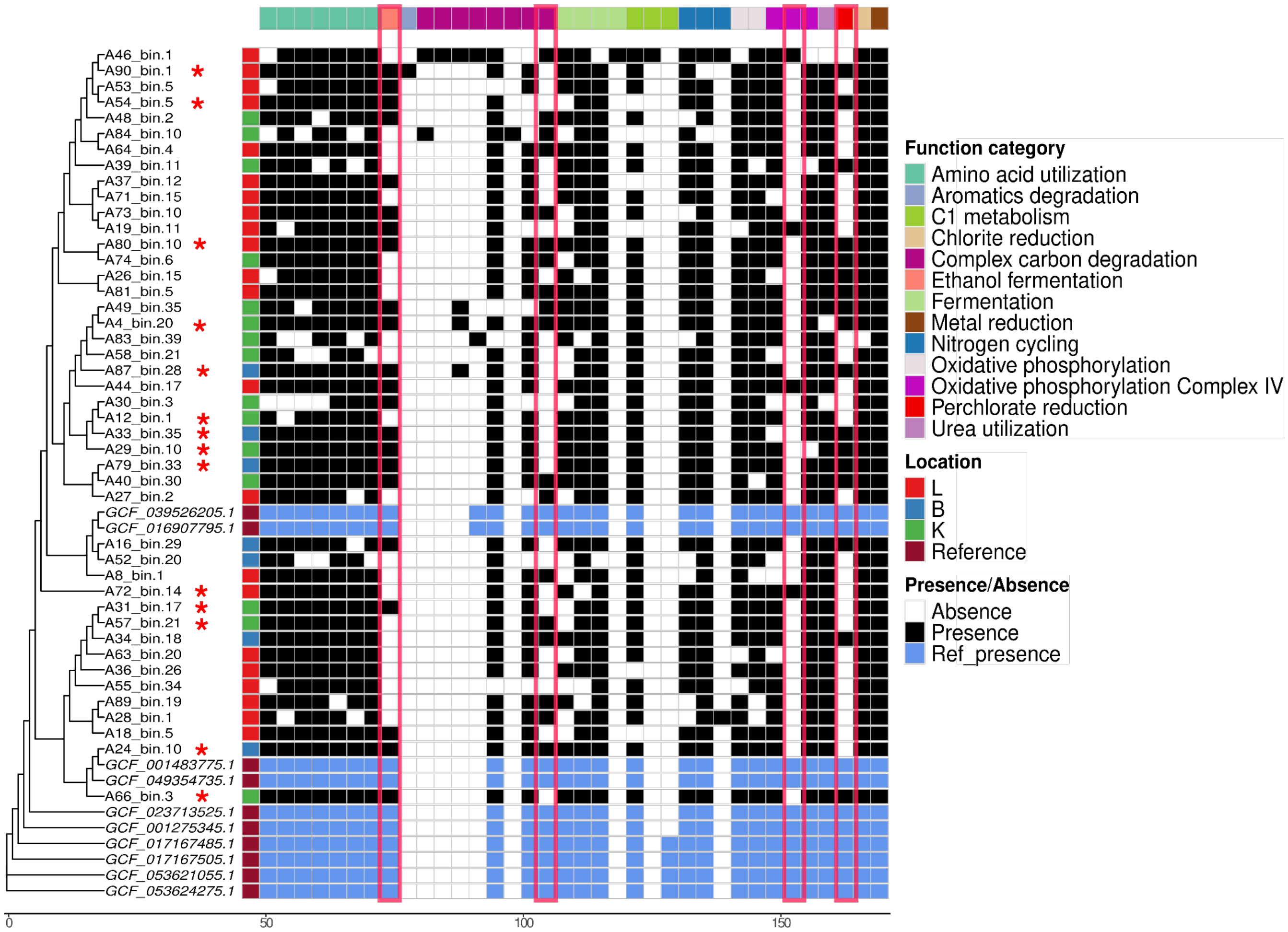


**Supplementary Fig. 6|Functional profile of *Kocuria palustris* MAGs and reference genomes.**
Presence/absence heatmap of annotated functional traits across *K. palustris* metagenome-assembled genomes (MAGs) and reference genomes, with hierarchical clustering based on functional similarity. Rows represent individual genomes and columns represent predicted functions grouped into broad metabolic categories, including amino acid utilization, aromatic compound degradation, C1 metabolism, chlorite reduction, complex carbon degradation, ethanol fermentation, metal reduction, nitrogen cycling, oxidative phosphorylation, oxidative phosphorylation complex IV, perchlorate reduction, and urea utilization. Black and white cells indicate presence and absence, respectively. The color bar at left denotes sample origin or genome source, distinguishing living area (L), bathroom (B), kitchen (K), and reference genomes. Red stars highlight the *K. palustris* MAGs recovered in this study with 90% completeness or more. The heatmap reveals conservation of core functional traits across genomes, alongside lineage-specific gain or loss of selected metabolic pathways, indicating functional diversification within this taxon across indoor house environments. The reference genomes had following accession numbers: GCF_001275345.1(culture contaminant), GCF_001483775.1 (Duodenal mucosa of Celiac disease patient), GCF_016907795.1(missing), GCF_017167485.1 (ISS environmental surface), GCF_017167505.1 (ISS environmental surface)GCF_023713525.1(Cleanroom),GCF_039526205.1 (rhizosphere of Typha angustiflora), GCF_049354735.1 (swimming pool water), GCF_053621055.1(ISS environmental surface), GCF_053624275.1 (ISS environmental surface).

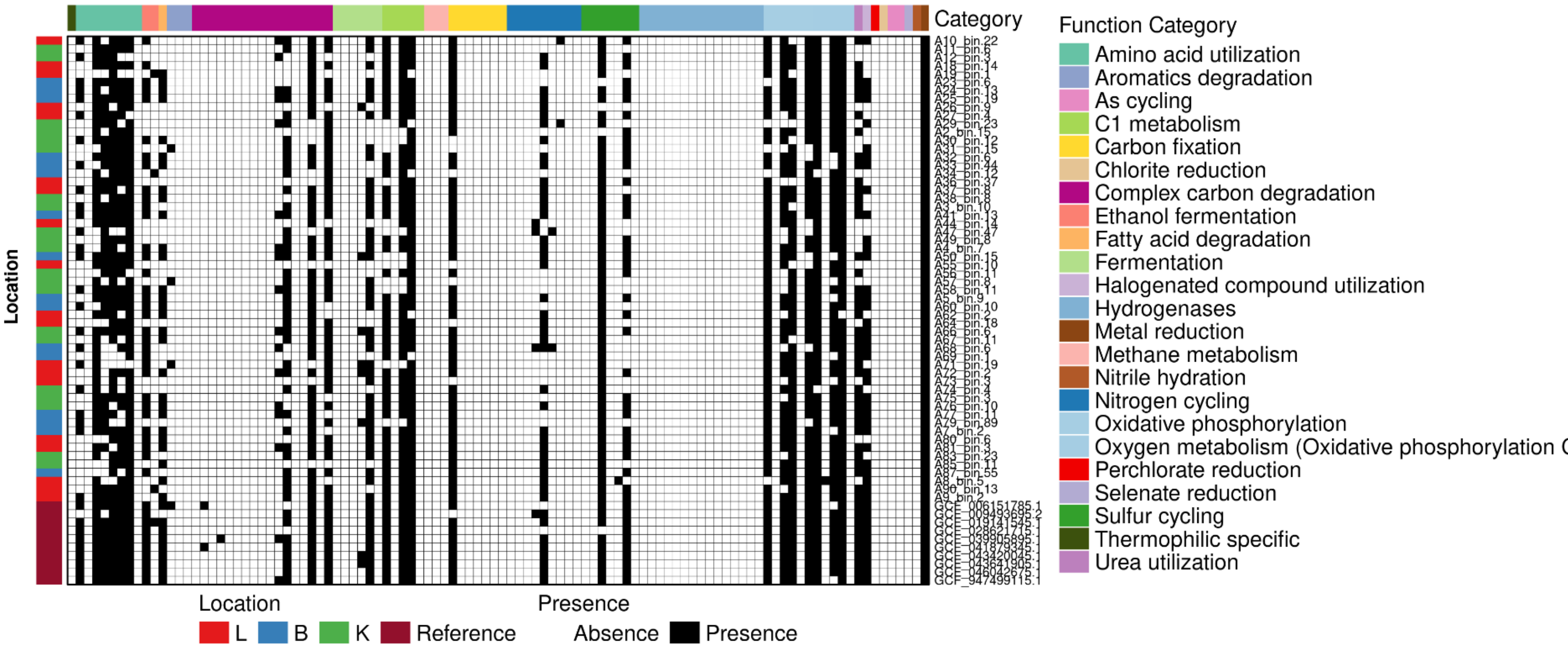


**Supplementary Fig. 7|Functional profile of *Paracoccus marcusii* MAGs and reference genomes.** Presence/absence heatmap of annotated functional traits across *Paracoccus marcusii* metagenome-assembled genomes (MAGs) and reference genomes. Rows represent individual genomes and columns represent predicted functions grouped into broad metabolic categories, including amino acid utilization, aromatic compound degradation, C1 metabolism, chlorite reduction, complex carbon degradation, ethanol fermentation, metal reduction, nitrogen cycling, oxidative phosphorylation, oxidative phosphorylation complex IV, perchlorate reduction, and urea utilization. Black and white cells indicate presence and absence, respectively. The color bar at left denotes sample origin or genome source, distinguishing living area (L), bathroom (B), kitchen (K), and reference genomes. Red stars highlight the *P. marcusii* MAGs recovered in this study with 90% or more completeness. The heatmap reveals conservation of core functional traits across genomes, alongside lineage-specific gain or loss of selected metabolic pathways, indicating functional diversification within this taxon across indoor house environments. The reference genomes had following accession numbers: GCF_006151785.1(sea water), GCF_009493695.2 (soil), GCF_019141545.1(carapace of Cancer pagurus), GCF_028621715.1(Halophyte), GCF_039905895.1(Suaeda maritima),GCF_041879345.1(Beaufort Sea), GCF_043420045.1(Sediments in Litoditis marina habitat), GCF_043641905.1(Sediments in Litoditis marina habitat), GCF_046042675.1(lake), GCF_947499115.1(phycosphere of marine macroalgae)

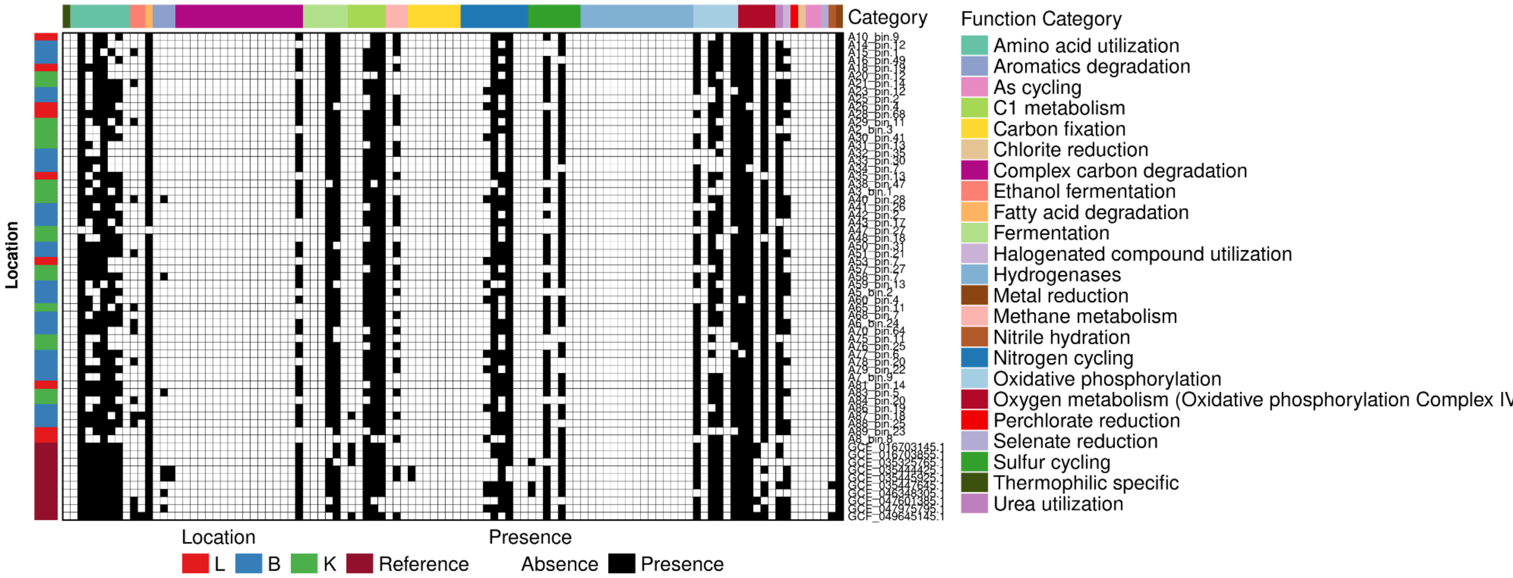


**Supplementary Fig. 8|Functional profile of *Ottowia* sp. 018060485 MAGs and reference genomes.** Presence/absence heatmap of annotated functional traits across *Paracoccus marcusii* metagenome-assembled genomes (MAGs) and reference genomes. Rows represent individual genomes and columns represent predicted functions grouped into broad metabolic categories, including amino acid utilization, aromatic compound degradation, C1 metabolism, chlorite reduction, complex carbon degradation, ethanol fermentation, metal reduction, nitrogen cycling, oxidative phosphorylation, oxidative phosphorylation complex IV, perchlorate reduction, and urea utilization. Black and white cells indicate presence and absence, respectively. The color bar at left denotes sample origin or genome source, distinguishing living area (L), bathroom (B), kitchen (K), and reference genomes. Red stars highlight the *P. marcusii* MAGs recovered in this study with 90% or more completeness. The heatmap reveals conservation of core functional traits across genomes, alongside lineage-specific gain or loss of selected metabolic pathways, indicating functional diversification within this taxon across indoor house environments.The reference genomes had following accession numbers: GCF_016703145.1 (activated sludge), GCF_016703855.1(activated sludge), GCF_035325765.1(full-scale industrial wastewater treatment plants sludge),GCF_035444425.1(Activated sludge and aerobic granule),GCF_035445925.1(Activated sewage sludge),GCF_035447645.1(Activated sludge of municipal wastewater treatment plants),GCF_046348305.1(full-scale hybrid biological wastewater treatment system),GCF_047601385.1(ant-built patch),GCF_047975795.1(crab intestines),GCF_049645145.1(Urban Sediment metagenome).

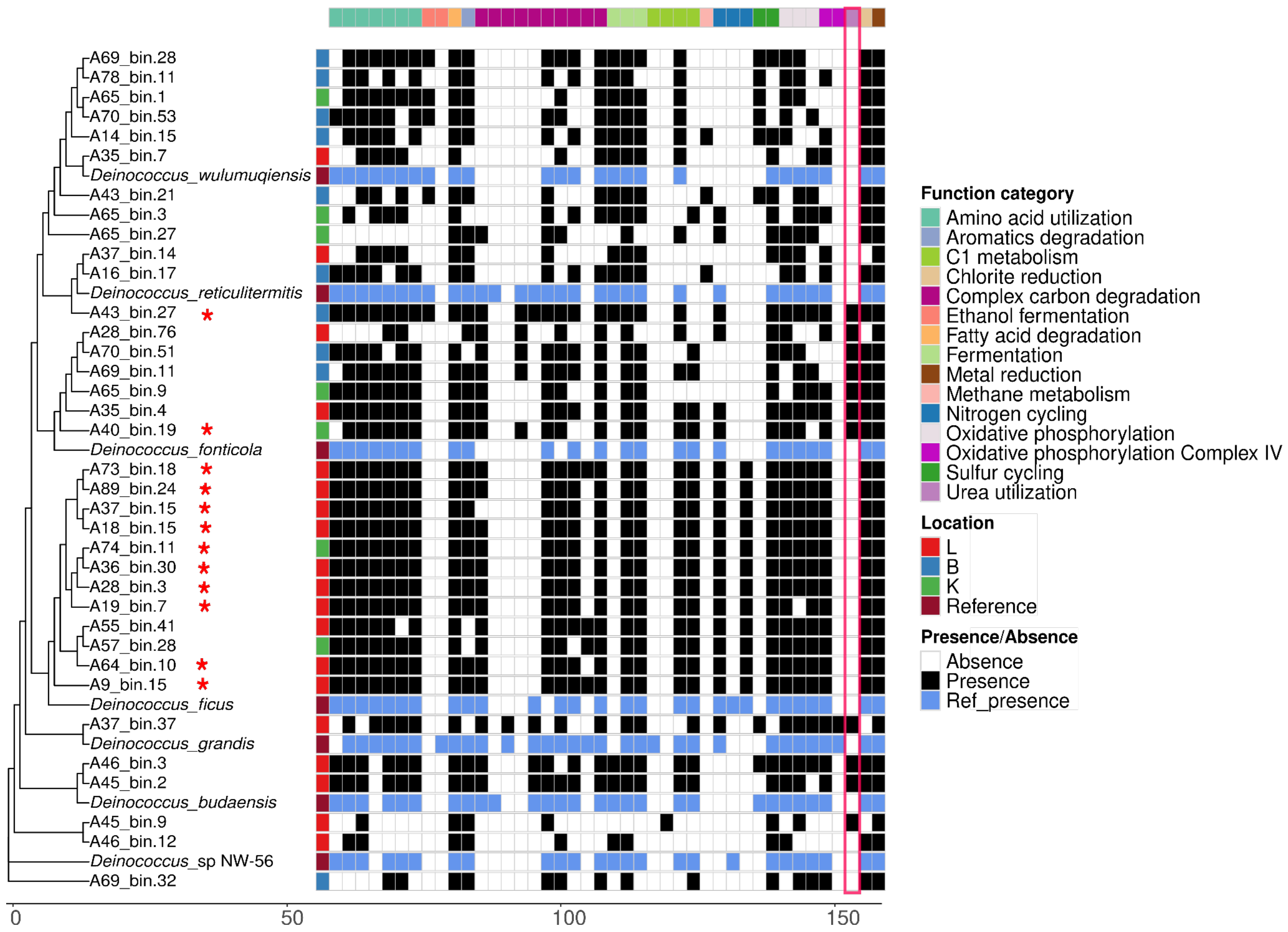


**Supplementary Fig. 9 | Functional repertoire of *Deinococcus* MAGs and reference genomes.** Presence/absence heatmap of annotated functional traits across *Deinococcus* metagenome-assembled genomes (MAGs) recovered in this study and reference genomes, with hierarchical clustering based on functional similarity. Rows represent individual genomes and columns represent predicted functions grouped into broad metabolic categories, including amino acid utilization, aromatic compound degradation, C1 metabolism, chlorite reduction, complex carbon degradation, ethanol fermentation, fatty acid degradation, fermentation, metal reduction, methane metabolism, nitrogen cycling, oxidative phosphorylation, oxidative phosphorylation complex IV, sulfur cycling, and urea utilization. Black and white cells indicate presence and absence, respectively. The left annotation bar denotes genome origin or location group (L, B, K, and reference genomes), and red stars highlight the *Deinococcus* MAGswith 90% and above completeness. The heatmap shows broad conservation of core metabolic features across genomes, together with lineage-specific differences in accessory functions, indicating functional diversification within this genus across indoor house environments. Reference genomes had the following accession numbers: *Deinococcus grandis* (freshwater): GCF_001485435, *Deinococcus* sp. NW-56 (water): GCF_002953415.1, *Deinococcus ficus* (rhizosphere) :GCA_003444775.1, *Deinococcus fonticola* (Diana-Hygieia radioactive thermal spring biofilm): GCF_004634215.1, Deinococcus *wulumuqiensis* R12 (soil):GCF_011067105.1, *Deinococcus budaensis*: GCF_014201885.1, *Deinococcus reticulitermitis*: GCF_900109185.1.

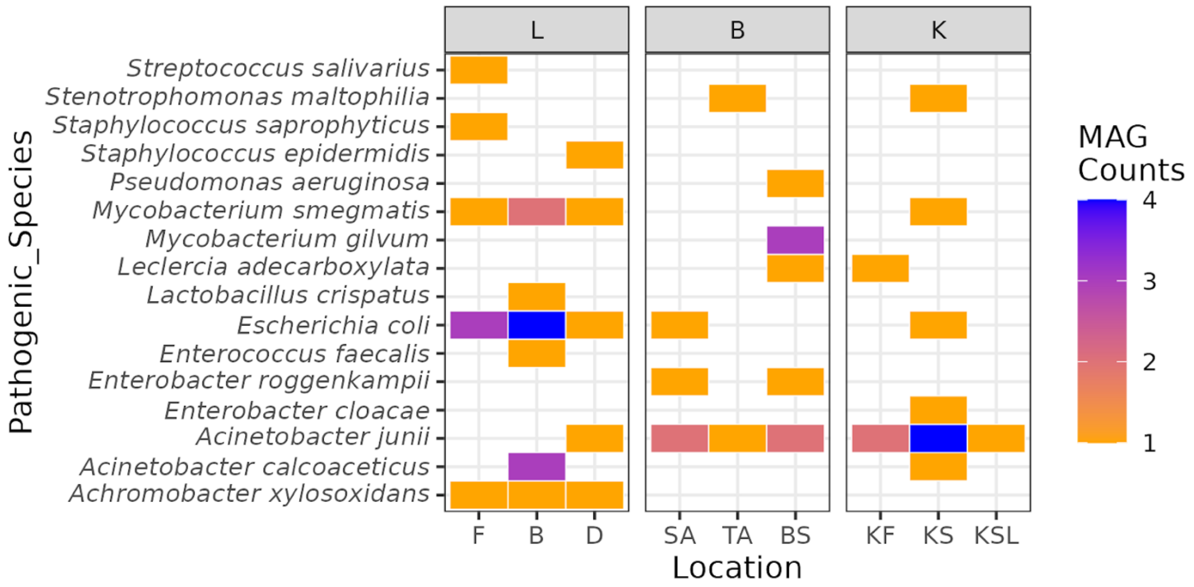


**Supplementary Fig. 10 | Distribution of pathogenic MAGs across house compartments and locations.**

Heatmap showing the distribution of pathogenic species-level metagenome-assembled genomes (MAGs) across living area (L), bathroom (B), and kitchen (K) compartments and their respective sampling locations. Rows represent pathogenic species and columns correspond to individual locations within each compartment. Color intensity indicates the number of MAGs recovered per species per location.

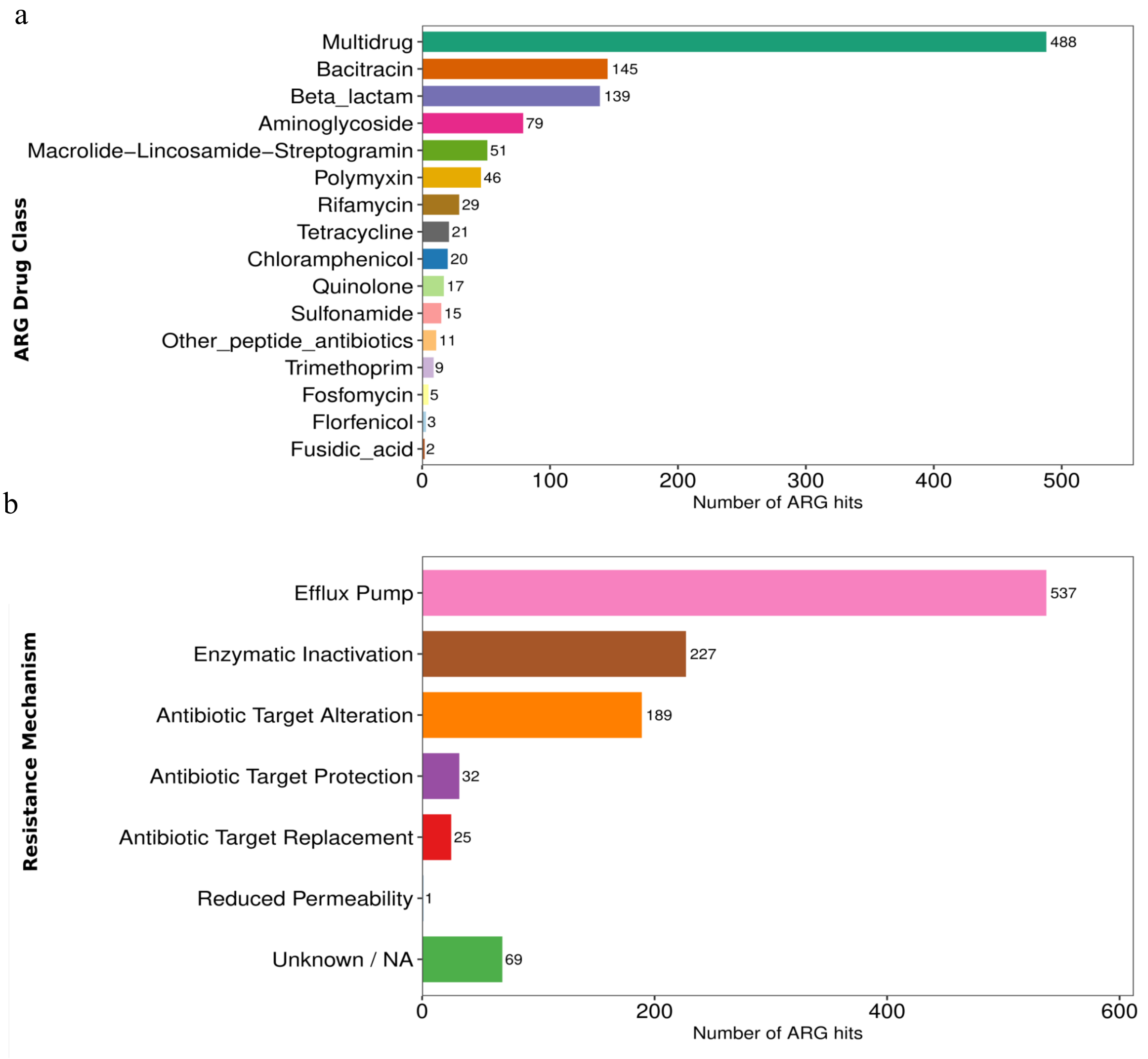


**Supplementary Fig. 11 | Distribution of the ARG hits**

(a)Number of ARG hits distributed to each drug class. (b) Distribution of the ARG hits by the resistance mechanism

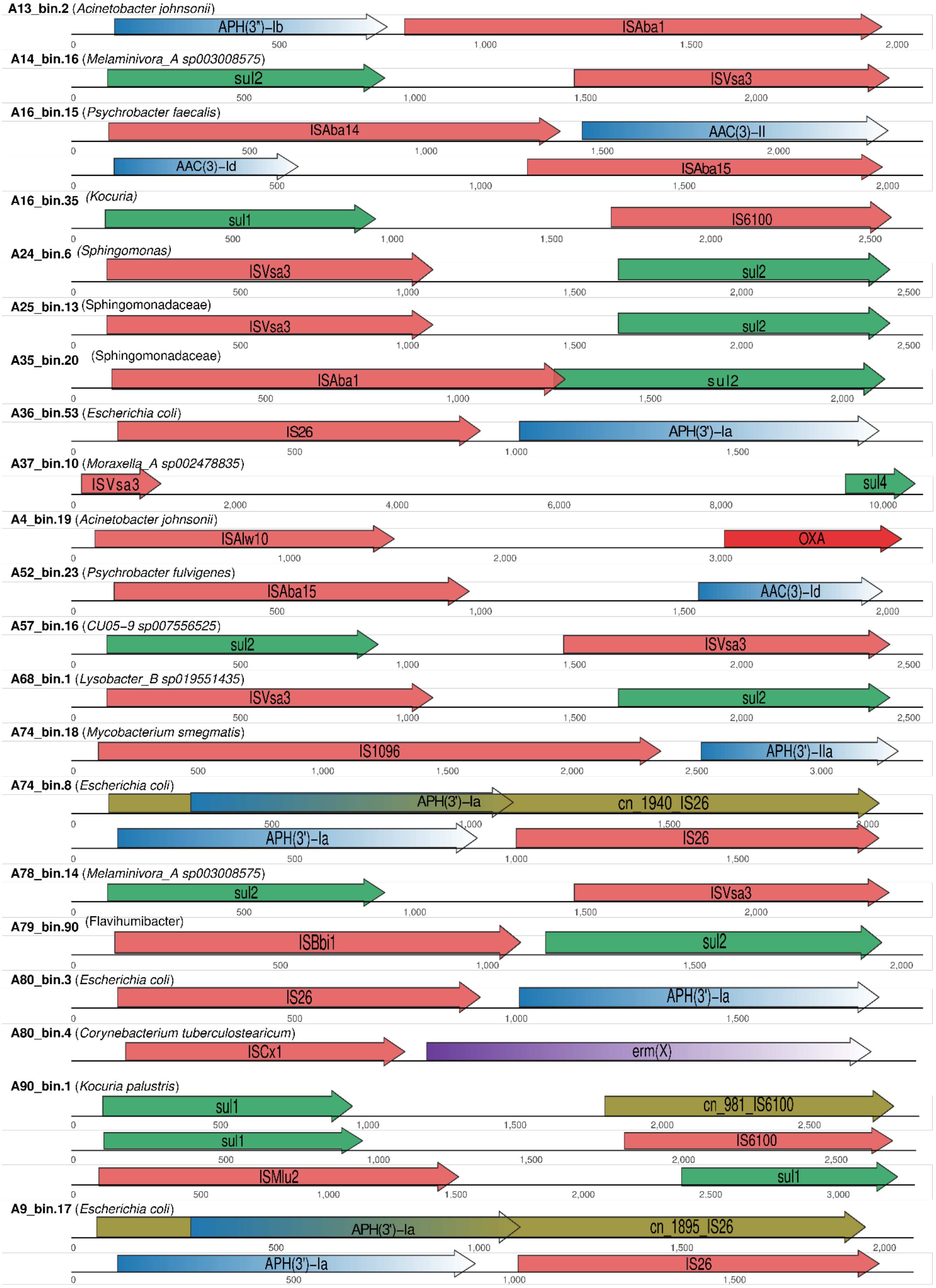
A13_bin.2 (Acinetobacter johnsonii)
APH(3")-Ib
ISAba1
A14_bin.16 (Melaminivora_A sp003008575)
sul2
ISVsa3
A16_bin.15 (Psychrobacter faecalis)
ISAba14
AAC(3)-II
AAC(3)-Id
ISAba15
A16_bin.35 (Kocuria)
sul1
IS6100
A24_bin.6 (Sphingomonas)
ISVsa3
sul2
A25_bin.13 (Sphingomonadaceae)
ISVsa3
sul2
A35_bin.20 (Sphingomonadaceae)
ISAba1
sul2
A36_bin.53 (Escherichia coli)
IS26
APH(3')-Ia
A37_bin.10 (Moraxella_A sp002478835)
ISVsa3
sul4
A4_bin.19 (Acinetobacter johnsonii)
ISAlw10
OXA
A52_bin.23 (Psychrobacter fulvigenes)
ISAba15
AAC(3)-Id
A57_bin.16 (CU05-9 sp007556525)
sul2
ISVsa3
A68_bin.1 (Lysobacter_B sp019551435)
ISVsa3
sul2
A74_bin.18 (Mycobacterium smegmatis)
IS1096
APH(3')-IIa
A74_bin.8 (Escherichia coli)
APH(3')-Ia
cn_1940_IS26
APH(3')-Ia
IS26
A78_bin.14 (Melaminivora_A sp003008575)
sul2
ISVsa3
A79_bin.90 (Flavihumibacter)
ISBbi1
sul2
A80_bin.3 (Escherichia coli)
IS26
APH(3')-Ia
A80_bin.4 (Corynebacterium tuberculostearicum)
ISCx1
erm(X)
A90_bin.1 (Kocuria palustris)
sul1
cn_981_IS6100
sul1
IS6100
ISMlu2
sul1
A9_bin.17 (Escherichia coli)
APH(3')-Ia
cn_1895_IS26
APH(3')-Ia
IS26

**Supplementary Fig. 12 | Detailed genomic layout of clinically relevant ARG-MGE pairs.** Schematic representation of clinically relevant antimicrobial resistance genes (ARGs) and mobile genetic elements (MGEs) located within a 10 kb distance of each other. Genomes are indicated by bold text and species names in italics. Arrows indicate the direction of transcription. Colors denote distinct functional classes of genes as indicated in the main text.
